# Pan-cancer *N*-glycoproteomic atlas of patient-derived xenografts uncovers FAT2 as a therapeutic target for head and neck cancers

**DOI:** 10.1101/2024.12.11.627962

**Authors:** Meinusha Govindarajan, Salvador Mejia-Guerrero, Shawn C Chafe, Shahbaz Khan, Wei Shi, Matthew Waas, Amanda Khoo, Lydia Y Liu, Vladimir Ignatchenko, Simona Principe, Lusia Sepiashvili, Nazanin Tatari, Chitra Venugopal, Petar Miletic, Maxwell Topley, Shan Grewal, Dillon McKenna, Maria-Jose Sandi, Nhu-An Pham, Alison Casey, Hyeyeon Kim, Christina Karamboulas, Jalna Meens, Peter Bergqvist, Begonia Silva, Patrick Chan, Liza Cerna-Portillo, Jasmine Chin, Abilasha Rao-Bhatia, Ming-Sound Tsao, Rama Khokha, Susie Su, Wei Xu, David Goldstein, Laurie Ailles, Vuk Stambolic, Fei-Fei Liu, Emma Cummins, Ismael Samudio, Sheila K Singh, Thomas Kislinger

**Author notes:** These authors contributed equally (Meinusha Govindarajan, Salvador Mejia-Guerrero).

## Abstract

Cell surface proteins offer significant cancer therapeutic potential attributable to their accessible membrane localization and central role in cellular signaling. Despite this, their promise remains largely untapped due to the technical challenges inherent to profiling cell surface proteins. Here, we employed *N*-glycoproteomics to analyze 85 patient-derived xenografts (PDX), constructing Glyco PDXplorer – an *in vivo* pan-cancer atlas of cancer-derived cell surface proteins. We developed a target discovery pipeline to prioritize proteins with favorable expression profiles for immunotherapeutic targeting and validated FAT2 as a head and neck squamous cancer (HNSC) enriched surface protein with limited expression in normal tissue. Functional studies revealed that FAT2 is essential for HNSC growth and adhesion through regulation of surface architecture and integrin-PI3K signaling. Chimeric antigen receptor (CAR) T cells targeting FAT2 demonstrated potent anti-tumor activity in HNSC models. This work lays the foundation for developing FAT2-targeted therapies and represents a pivotal resource to inform therapeutic target discovery for multiple cancers.

**HIGHLIGHTS:**

- Pan-cancer landscape of cancer-derived cell surface proteins detected *in vivo*
- Development of a multi-omic discovery pipeline to prioritize proteins with optimal expression profiles as immunotherapy targets
- Identification and validation of FAT2 as a head and neck squamous cancer enriched surface protein with minimal expression in normal tissues
- FAT2 coordinates cell surface organization, adhesion, growth and survival through the integrin-PI3K-AKT pathway
- FAT2 CAR T cells demonstrate anti-tumour activity in pre-clinical models

## INTRODUCTION

Targeted therapies have been recognized as one of the most pivotal advancements towards reducing cancer-related mortality worldwide^1^. In contrast to conventional cancer therapies, which damage both cancer and normal cells, thereby incurring toxicity to healthy tissues, targeted therapies aim to direct their toxicity specifically to cancer cells, sparing normal tissues^2^. Cell surface proteins represent attractive therapeutic targets due to their accessible subcellular localization and their involvement in essential signaling pathways, often dysregulated in cancer^3,4^. Unlike intracellular proteins, cell surface antigens can also be targeted by a myriad of antibody-based approaches including Fc-mediated therapeutics, antibody-drug conjugates (ADCs), single-chain variable fragment (scFv)-based formats (*i.e.* BiTEs, DARTs, single domains, TandAbs, crossmabs), and chimeric antigen receptor (CAR) T cells^4^. Notably, cell surface proteins constitute over 60% of FDA-approved protein drug targets underscoring their value as therapeutic targets^5^. Thus, targeted therapeutics tailored specifically for cancer cells based on surface marker expression have tremendous potential to decrease tumor burden with minimal drug-toxicity and could offer attractive alternatives to current standard of care treatments.

Despite their importance as drug targets, only a few cell surface proteins with established roles in cancer biology are targeted by current therapeutic antibodies^6^. Therefore, the full potential of the cell surface proteome (*i.e.* surfaceome) for cancer therapeutic targeting has yet to be realized. A major obstacle for the detection of novel cancer specific surface antigens are the challenges associated with comprehensively defining the surfaceome. Most surface antigen discovery efforts are based on the use of transcriptomic data to infer surface protein expression^7^. However, RNA abundance does not always correlate with protein abundance and transcriptomics data cannot empirically determine protein subcellular localization^8,9^. Although target discovery platforms based on mass spectrometry (MS)-based proteomics quantify thousands of proteins in a single sample^9^, cell surface proteins are typically under-represented in such global proteomics datasets due to their low abundance and biophysical properties^4^. Hence, enrichment prior to MS analysis is warranted for extensive surface protein profiling. As over 80% of cell surface and secreted proteins are *N*-glycosylated^10,11^, specific chemoproteomic enrichment of *N*-glycoproteins is a powerful tool to elucidate the surfaceome.

Patient-derived xenografts (PDX) are generated by engrafting human tumors into immunocompromised mice, enabling *in vivo* interrogation of patient tumors. PDXs faithfully recapitulate histopathological, molecular, and clinical features of their matched patient tumors rendering these models highly conducive for cancer discovery and validation studies^12^. Here, we applied *N*-glycoproteomics to 85 unique PDX models spanning seven tumor types and generated Glyco PDXplorer – an *in vivo* pan-cancer atlas of cancer-derived surface proteins. We devised a multi-omic cell surface target discovery pipeline and prioritized 290 proteins as targeted therapy candidates and validated FAT2 as a previously undescribed head and neck squamous cancer (HNSC)-enriched cell surface protein with limited expression in normal tissue. *In vitro* and *in vivo* knockdown of FAT2 uncovered its essential function in adhesion, growth and survival, mediated through control of cell surface organization and the integrin-PI3K-AKT signaling axis in HNSC. Finally, our proof-of-concept engineering and evaluation of a FAT2 CAR T construct highlights the potential of FAT2 as a novel immunotherapy target in HNSC and validates our platform for target discovery. The pan-cancer *N*-glycoproteomic PDX data presented in this work has been integrated into an interactive data portal (http://kislingerlab.uhnres.utoronto.ca/glycoPDXplorer/) to accelerate the development of targeted therapies for additional cancers.

## RESULTS

### Pan-cancer PDX *N*-glycoproteome atlas of cancer-derived surface proteins

PDX tumors contain human cancer cells and murine tumor microenvironmental features (*e.g.* fibroblasts, vasculature, limited immune populations *etc.*)^12,13^. Species based deconvolution of PDX molecular profiles is therefore a simple and powerful approach to discriminate between cancer and tumor microenvironment (TME) in bulk tissue samples. Notably, *in silico* species comparison of theoretical *N*-glycopeptides and global tryptic peptides revealed that *N*-glycopeptides are significantly less conserved between human and mouse proteomes compared to global tryptic peptides **(Figures S1A & S1B**), highlighting the advantage of using *N*-glycoproteomics for species discrimination. Chemoproteomic enrichment of *N*-glycoproteins in PDX tumors thus represents an intriguing strategy to determine cancer cell-derived surface proteins *in vivo*. Here, we applied a hydrazide-based *N*-glycoproteomic enrichment method, *N*-glycocapture^14–16^, to profile 85 unique PDX models spanning seven tumor types (glioblastoma [GBM], HNSC, lung squamous carcinoma [LUSC], lung adenocarcinoma [LUAD], pancreatic adenocarcinoma [PAAD], colorectal adenocarcinoma [COAD] and high-grade serous ovarian carcinoma [OV]^14^) and three distinct PDX engraftment sites (intracranial, subcutaneous and intraperitoneal) (**Figure 1A**, **Table S1**). *N*-glycoproteome profiling of PDX models was highly reproducible as evident by minimal variability between engraftment, processing and technical replicates (**Figure S1C**). In total, 6,378 *N*-glycopeptides mapping to 2,873 *N*-glycoproteins were detected (**Table S1**). Over 90% of detected proteins were predicted to have a surface membrane localization and/or contain a signal peptide (**Figure 1B**)^10^, underscoring the specificity of *N*-glycoproteomics for enrichment of cell surface and secreted proteins.

**Figure 1.**
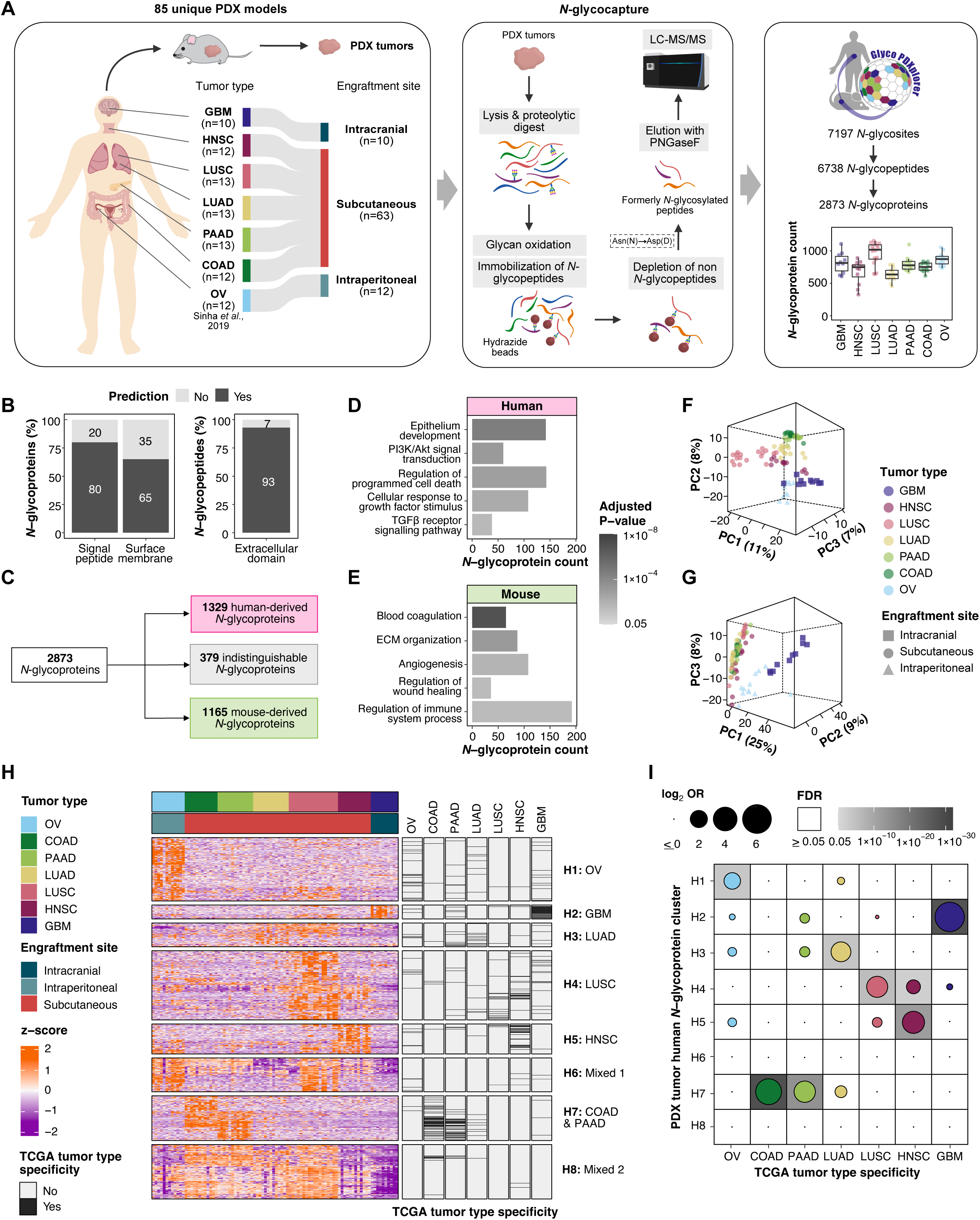
Pan-cancer PDX *N*-glycoproteome atlas of cancer-derived surface proteins. **(A)** Overview of patient-derived xenograft (PDX) cohort (n=85) and workflow for *N*-glycoprotein enrichment used to generate Glyco PDXplorer. Tumor types are indicated by official study abbreviations from The Cancer Genome Atlas (TCGA). **(B)** Percentage of total *N*-glycoproteins (left) and *N*-glycopeptides (right) with predicted surface and secreted protein annotations^10^. **(C)** Species breakdown of detected *N*-glycoproteins. **(D & E)** Select uniquely enriched GO: Biological Processes in **(D)** human-derived PDX *N*-glycoproteins and **(E)** mouse-derived PDX *N*-glycoproteins. **(F & G)** Three-dimensional principal component analysis of PDX tumors using **(F)** human-derived *N*-glycoproteins and **(G)** mouse-derived *N*-glycoproteins. Color indicates tumor type and shape denotes engraftment site. **(H)** Expression heatmap of human PDX *N*-glycoproteins that are differentially expressed by tumor type (ANOVA, False Discovery Rate [FDR] < 0.05, n=857 *N*-glycoproteins). *N*-glycoproteins (rows) are split by Consensus clustering. Covariate bars indicate tumor type specificity in patient tumors as determined from TCGA data^17^. **(I)** Dot plot depicting over-representation of TCGA tumor type specific gene-products in respective human PDX *N*-glycoprotein clusters. Dot size indicates the enrichment odds ratio (OR), and background shading represents FDR. See also **Figure S1**.

Following stringent species assignment (see **Methods**), 1329 *N*-glycoproteins were defined as human-derived, and 1165 proteins were considered mouse-derived (**Figures 1C and S1D**). Differences in species composition among tumor types were minimal except for GBM samples which comprised mostly mouse *N*-glycopeptides (**Figure S1E**). This is likely due to the infiltrative and diffusive nature of intracranial GBM tumors and the consequent sample processing of entire resected brains rather than micro-dissected tumors. Detected human PDX proteins were uniquely enriched in cancer cell processes including epithelial development and cell death regulation (**Figure 1D**) whereas detected mouse PDX proteins were uniquely enriched in TME processes related to extracellular-matrix (ECM) and vasculature (**Figure 1E**, **Table S1**). Principal component analysis revealed that PDX tumors separated by tumor type when analyzing human-derived proteins (**Figure 1F**), and in contrast, clustered by engraftment site when considering mouse-derived proteins (**Figure 1G**). These data reinforce the ability of bioinformatic species assignment to distinguish between cancer-derived (*i.e.* human) and TME-derived (*i.e.* mouse) proteins in PDX tumors.

To gain insights into cancer cell differences between tumor types, consensus clustering was performed on human PDX proteins differentially expressed between tumor types. This analysis uncovered eight robust *N*-glycoprotein clusters – mostly enriched in distinct tumor types (**Figure 1H**). Pathway analysis indicate that human *N*-glycoprotein clusters reflect unique biological processes (**Figure S1F**, **Table S1**). For example, proteins in H1, the OV cluster, were enriched in metabolic processes and anatomical structure development, whereas proteins in H2, the GBM cluster, were associated with neuronal processes. To evaluate whether the differences among PDX tumor types recapitulate patient tumor type differences, we compared our findings to tumor type specific transcripts and proteins determined using patient data (*i.e.* The Cancer Genome Atlas [TCGA]^17^ and Clinical Proteomic Tumor Analysis Consortium [CPTAC]^18^ data) (**Figures S1G & S1H**). Indeed, patient tumor type specific transcripts and proteins were concordant with PDX human *N*-glycoprotein clusters (**Figures 1H, 1I & S1I**). These findings indicate that patient tumor type differences are preserved in PDX models. Together, the multi-cancer human PDX *N*-glycoproteomics data generated in this study – from here on referred to as Glyco PDXplorer – represents a comprehensive *in vivo* resource of cancer-derived surface proteins.

### Prioritization and evaluation of cancer enriched surface proteins with limited normal expression as therapeutic target candidates

To evaluate the value of the Glyco PDXplorer atlas for therapeutic target discovery, we first assessed the number of clinical (*i.e.* FDA approved and/or in clinical trial) solid tumor immunotherapy (*i.e.* ADC and CAR T) targets detected in the dataset. Over 60% of clinical immunotherapy targets for solid tumors, irrespective of specific tumor type, were detected in our study (**Figure 2A**). Furthermore, the surface proteins detected in Glyco PDXplorer were significantly enriched in clinical targets compared to the remaining human surfaceome (**Figures 2B, S2A & S2B**). These analyses demonstrate that Glyco PDXplorer is advantageous for detecting clinically relevant proteins and highlight its potential for target discovery.

**Figure 2:**
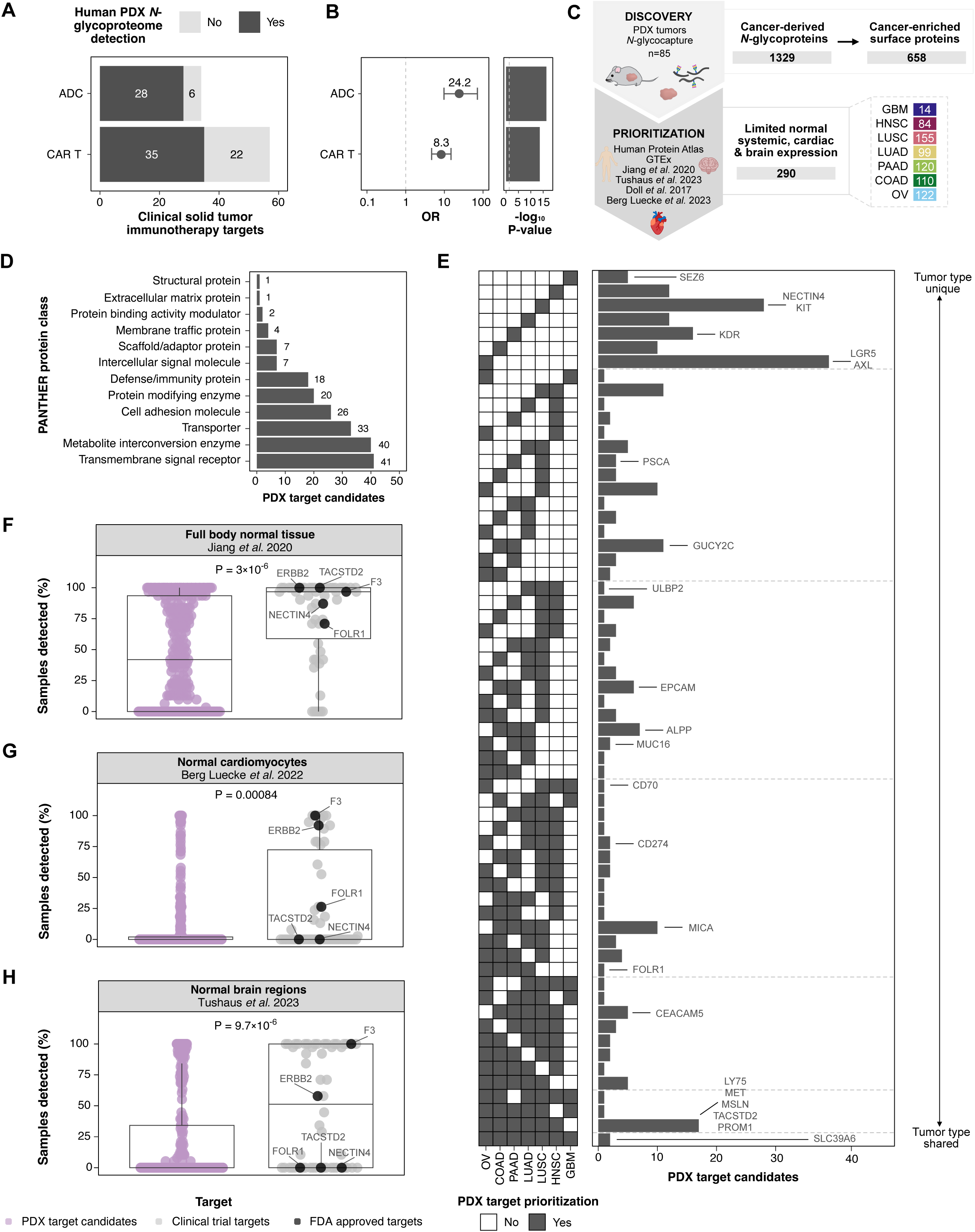
Prioritization and evaluation of cancer-enriched surface proteins with limited normal expression as therapeutic target candidates. **(A)** Number of solid tumor clinical (*i.e*. FDA approved and/or under clinical trial) immunotherapy (*i.e.* antibody drug conjugate [ADC] and/or chimeric antigen receptor [CAR] T cell therapy) targets detected in the human Glyco PDXplorer data. **(B)** ORs (left) and P-values (right, Fisher’s exact test) demonstrating the likelihood of detecting a clinical immunotherapy target in the human PDX *N*-glycoproteome atlas against all other surface proteins in the human proteome. **(C)** PDX driven strategy for prioritization of cancer enriched surface proteins with limited expression in normal tissue (*i.e.* PDX target candidates). Grey boxes indicate number of unique proteins across entire dataset for each stage. Final PDX target candidate counts for individual tumor types are indicated in colored boxes. **(D)** PANTHER protein classification of PDX target candidates. **(E)** Upset plot depicting overlap of PDX target candidates between tumor types. Clinical immunotherapy targets are labelled on respective intersections for which they were prioritized as putative targets by the PDX pipeline. **(F-H)** Frequency of sample detection of PDX target candidates (purple) *vs* all solid tumor clinical targets (grey) in normal tissues profiled by **(F)** Jiang *et al*., 2020^20^ **(G)** Berg Luecke *et al.,* 2022^23^ and **(H)** Tushaus *et al.,* 2023^21^. FDA approved immunotherapy targets are annotated in black. P-values from unpaired Mann-Whitney U tests. See also **Figure S2**.

An ideal immunotherapy target ought to: (1) be highly expressed on the surface of cancer cells for potent targeting; and (2) have limited expression in normal cells to minimize systemic toxicities. Guided by these principles, we devised a two-stage PDX discovery pipeline and applied it individually to the seven tumor types (**Figure 2C**). First, we filtered the human (*i.e.* cancer-derived) Glyco PDXplorer data for surface proteins highly abundant in PDX tumors from the respective tumor type (see **Methods**), resulting in the identification of 658 unique surface proteins enriched in at least one of seven tumor types. The second stage integrated published multi-omic normal tissue data^5,19–24^ (**Table S2**) to prioritize proteins with limited abundance across all normal tissues and specifically in vital organs (*i.e.* heart and brain) to reduce the likelihood of normal tissue toxicity (**Figure S2C**). Two hundred and ninety unique cell surface proteins highly abundant in at least one of the seven tumor types with restricted expression in normal tissue were prioritized as putative therapeutic targets – from here on referred to as PDX target candidates (**Figure 2C**). Consistent with their surface localization, PDX target candidates were comprised predominantly of signal receptors, transporters and adhesion molecules (**Figure 2D**). Many PDX target candidates (122/290) were only nominated for a single tumor type, yet 21 proteins were prioritized as potential targets for at least 6/7 tumor types (**Figure 2E**). These multi-cancer PDX target candidates include known targets such as TACSTD2 – a FDA approved ADC target for breast and urothelial cancers^25^, and SLC39A6 – a clinical trial ADC target for breast cancer^26^. Additionally, FOLR1 – an FDA approved ADC target for OV^25^ – was prioritized by our pipeline as a potential target for OV and three additional tumor types (**Figure 2E**), exemplifying the use of our PDX driven target discovery pipeline to nominate additional cancer type indications for existing targets. Finally, to benchmark the PDX driven target discovery pipeline, we compared the expression profiles between PDX target candidates and clinical immunotherapy targets across 12 normal tissue data metrics^5,19–24^. These extensive, multi-omic analyses revealed that PDX target candidates exhibit lower or comparable normal tissue expression as current clinical targets (**Figures 2F-2H, S2D-S2F**). Together, these findings underscore the potential of PDX *N*-glycoproteomics as a robust platform for identifying proteins with optimal expression profiles for immunotherapeutic targeting.

### Identification and validation of FAT2 as a novel HNSC enriched surface protein with limited expression in normal tissue

As a proof-of-principle demonstration of our PDX *N*-glycoproteomic pipeline for the discovery of previously undescribed targets, we sought to identify and validate a novel target candidate for HNSC – a tumor type we have previously worked with^27–30^. To narrow down the 84 PDX target candidates prioritized for HNSC (**Figure 2C**) and select a single protein for detailed interrogation, we conducted additional *N*-glycoproteomic profiling for large-scale verification of HNSC patient tumor expression and surface localization, respectively (**Figure 3A, Tables S3 and S4**). As we recognize that the lack of immune system in PDX models may impact cancer cell expression profiles compared to patient tumors, we performed *N*-glycocapture on 87 HNSC patient tumor specimens to corroborate patient expression of target candidates (**Table S3**). To validate cell surface expression of candidates in HNSC, we also utilized Cell Surface Capture (CSC)^31^ – an alternative hydrazide-based *N*-glycoproteomic method that provides empirical evidence of cell surface localization – in a panel of nine established cell lines (6 HNSC, 2 additional non-HNSC squamous cell carcinoma [SCC] and one normal oral epithelial [NOE] cell lines) (**Figure 3B, Table S4**). Integration of these two HNSC *N*-glycoproteomic datasets with our PDX pipeline retained 31 high-confidence candidates (**Figure 3A**) which we further ranked based on: (1) frequency of detection and expression in HNSC PDX; (2) frequency of detection and expression in HNSC patient tumors; and (3) differential surface expression between SCC cells and the NOE cell controls (**Figure 3C**). Protocadherin FAT2, a relatively unexplored member of the cadherin family of adhesion molecules^32,33^, emerged as the top ranked candidate and was selected for investigation. Intriguingly, FAT2 exhibited differential surface abundance between HNSC cell lines and NOE controls comparable to that of EGFR, which is the only FDA-approved antibody target currently available for HNSC (**Figure S3A**). Immunofluorescence microscopy provided orthogonal evidence of a cell surface localization of FAT2 (**Figure S3B**). Increased FAT2 protein in HNSC cell lines compared to control NOE cells was substantiated with immunoblotting using both a commercial (Santa Cruz Biotechnologies [SCB]) and a newly generated anti-human FAT2 antibody (A088) (**Figure S3C**). Detailed antibody characterization revealed that A088 has high affinity and specificity to the extracellular domain of FAT2, thus expanding the molecular toolbox available to interrogate FAT2 (see **Methods**, **Note S1** and **Figures S3D-S3L**).

**Figure 3.**
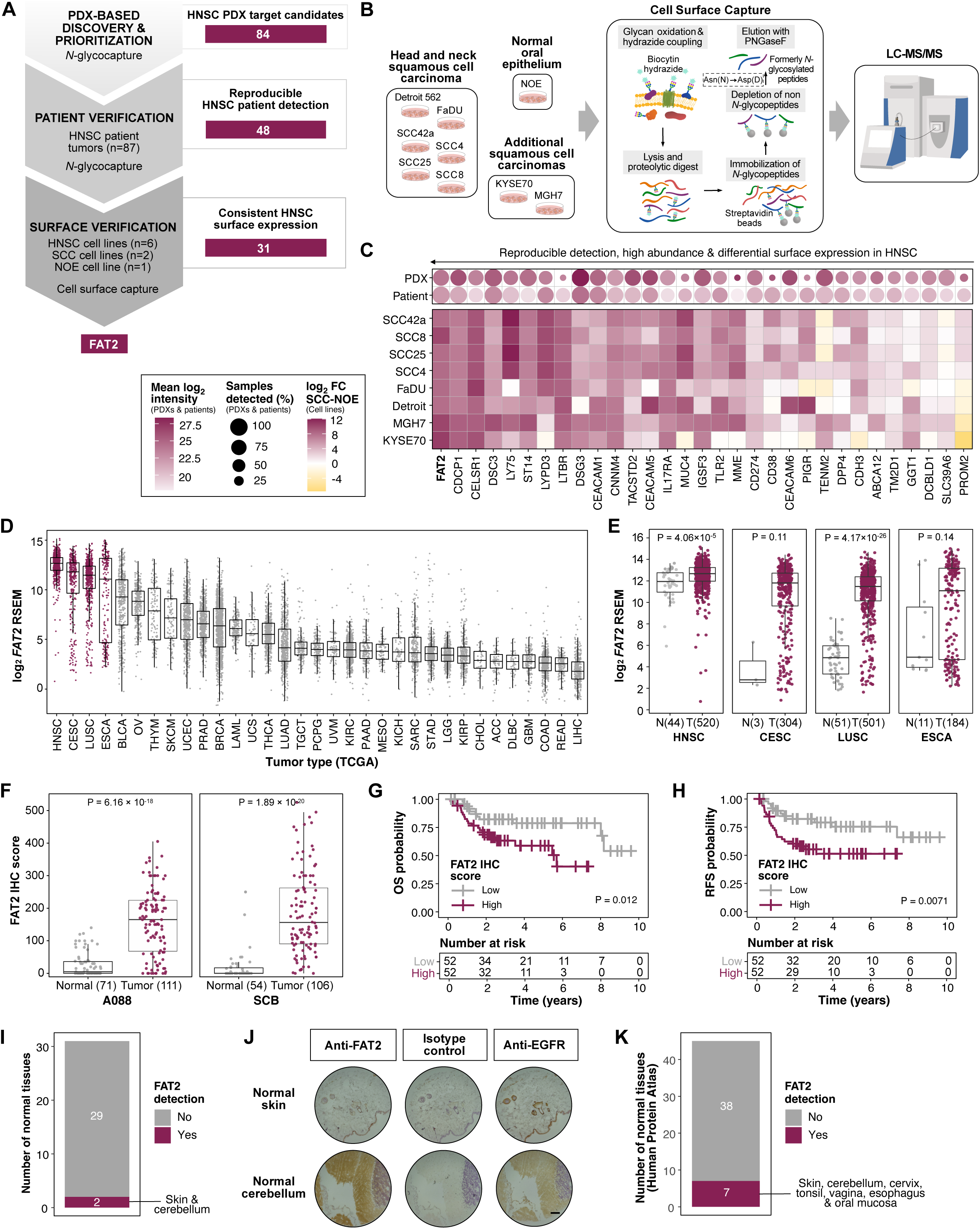
Identification and validation of FAT2 as a HNSC enriched surface protein with limited expression in normal tissue. **(A)** Expansion of the head and neck squamous cell carcinoma (HNSC) PDX *N*-glycoproteomic target discovery strategy from Figure 2C with *N*-glycoproteomic verification of HNSC patient detection and cell surface expression. **(B)** Overview of cell surface capture (CSC) *N*-glycoproteomic approach used to verify HNSC cell surface expression in **(A)**. Three replicates per cell line. **(C)** Thirty-one high confidence HNSC PDX target candidates ordered by detection and abundance in HNSC PDX (n=12) and HNSC patients (n=87) (top) and differential surface expression between squamous cell carcinoma (SCC) cell lines (n=8) and normal oral epithelial (NOE) cells (n=1) (bottom). **(D)** *FAT2* transcript expression in primary human tumors profiled by TCGA^17^. Tumor types use official TCGA Study Abbreviations. SCC tumors are colored in burgundy. **(E)** *FAT2* transcript expression in SCC primary human tumors (T) and adjacent normal tissue (N) from TCGA. Sample number indicated in parentheses and statistical differences were determined with an unpaired Student’s t-test. **(F)** Immunohistochemistry (IHC) evaluation of FAT2 in an in-house tissue microarray (TMA) containing HNSC tumor cores and adjacent normal tissue. Left panel: Newly developed anti-FAT2 antibody (A088). Right panel: commercial antibody (Santa Cruz Biotechnologies [SCB]). Sample number indicated in parentheses and statistical differences were assessed with an unpaired Student’s t-test. **(G & H)** HNSC patient **(G)** overall survival (OS) and **(H)** recurrence free survival (RFS) probability based on median dichotomized FAT2 expression as stained by SCB antibody. Statistical significance was calculated with a log-rank test. **(I)** Number of normal tissues stained positive for FAT2 by IHC using the anti-FAT2 antibody A088 with a commercially available TMA. **(J)** Representative images of positive FAT2 IHC detection in normal skin and cerebellum tissue as indicated in **(I).** Isotype control (IgG2a) and anti-epidermal growth factor receptor (EGFR) antibodies were used as negative and positive controls, respectively. Scale bar = 200 μm. **(K)** Number of normal tissues stained positive for FAT2 by IHC as per Human Protein Atlas^5^. See also **Figures S3 and S4**.

To evaluate FAT2 expression in other cancer types, we first used our multi-cancer PDX resource (**Figure 1A**). FAT2 was robustly detected in both HNSC and LUSC tumors – the only two squamous cancer types profiled in this study (**Figure S3M**). Similar analyses of TCGA mRNA expression data^17^ uncovered that the highest median expression of *FAT2* was observed in HNSC followed by three other tumor types with squamous cell origin (cervix squamous cell carcinoma [CESC], LUSC and esophageal squamous cell carcinoma [ESCA]), suggesting that FAT2 is a squamous cancer enriched cell surface marker (**Figure 3D**). Further interrogation of TCGA data indicated that *FAT2* was upregulated in tumor tissues, compared to adjacent normal tissues, in all four squamous tumor types (**Figure 3E**). To validate these findings at the protein level, we utilized a richly annotated in-house HNSC tissue microarray (TMA) that contains both tumor and adjacent normal tissue cores (n=117, **Table S5**). Independent immunohistochemistry (IHC) assessments with both SCB and A088 FAT2 antibodies indicated higher abundance of FAT2 in tumors compared to adjacent normal tissues (**Figure 3F**). Notably, increased FAT2 protein was associated with significantly worse overall survival and recurrence free survival probabilities (**Figures 3G, 3H**, **S3N, S3O**; **Table S5**), using both antibodies independently.

To verify the limited systemic normal tissue expression of FAT2, we performed IHC with the A088 FAT2 antibody on a commercially available TMA of 31 normal tissues. Strikingly, FAT2 was only observed in tissue cores from skin and cerebellum while the remaining 29 analyzed tissues displayed no detectable FAT2 signal (**Figures 3I, 3J & S4A; Table S5**). This corroborated Human Protein Atlas IHC data^5^ in which FAT2 was detected in merely 7/45 normal tissues, including cerebellum and skin (**Figure 3K**). We analyzed spatially and cell-type resolved datasets of normal skin^34^ and brain^35^ to gain additional insights into FAT2 abundance in these tissue types. FAT2 was the most abundant in the epidermis (**Figure S4B**), the layer of skin with no vasculature^36^ and thus, less accessible to blood-based delivery of therapeutics. Interestingly, FAT2 was detected in normal skin at lower levels than EGFR (**Figure S4B**) – a HNSC antibody target associated with noteworthy skin toxicities^37^. In the brain^35^, *FAT2* expression was predominately restricted to a subset of lower and upper rhombic lip cells, the precursors of the cerebellum (**Figure S4C**). Together, these data validate FAT2 – a novel PDX target candidate – as a HNSC enriched surface protein with limited normal tissue expression in humans.

### FAT2 is essential for HNSC growth *in vitro* and *in vivo*

While preliminary reports implicate a potential association between FAT2 and squamous cancers^38–41^, a comprehensive characterization of FAT2 expression and function in HNSC remains elusive. To evaluate the significance of elevated FAT2, siRNAs were used to transiently downregulate FAT2 expression in a panel of eight cell lines (6 HNSC and 2 non-HNSC SCC) (**Figure S5A**). Using three independent siRNAs, knockdown of FAT2 substantially reduced cellular proliferation and colony-forming capacity of all tested FAT2-positive cancer cell lines compared to a non-targeting scramble siRNA (**Figures 4A, 4B, S5B**). In contrast, treatment of FAT2-negative cell lines, NOE and HT29, with FAT2 siRNAs had no effect on their proliferation (**Figure 4A)**. These results indicate that FAT2 is required for HNSC growth and survival *in vitro*.

**Figure 4.**
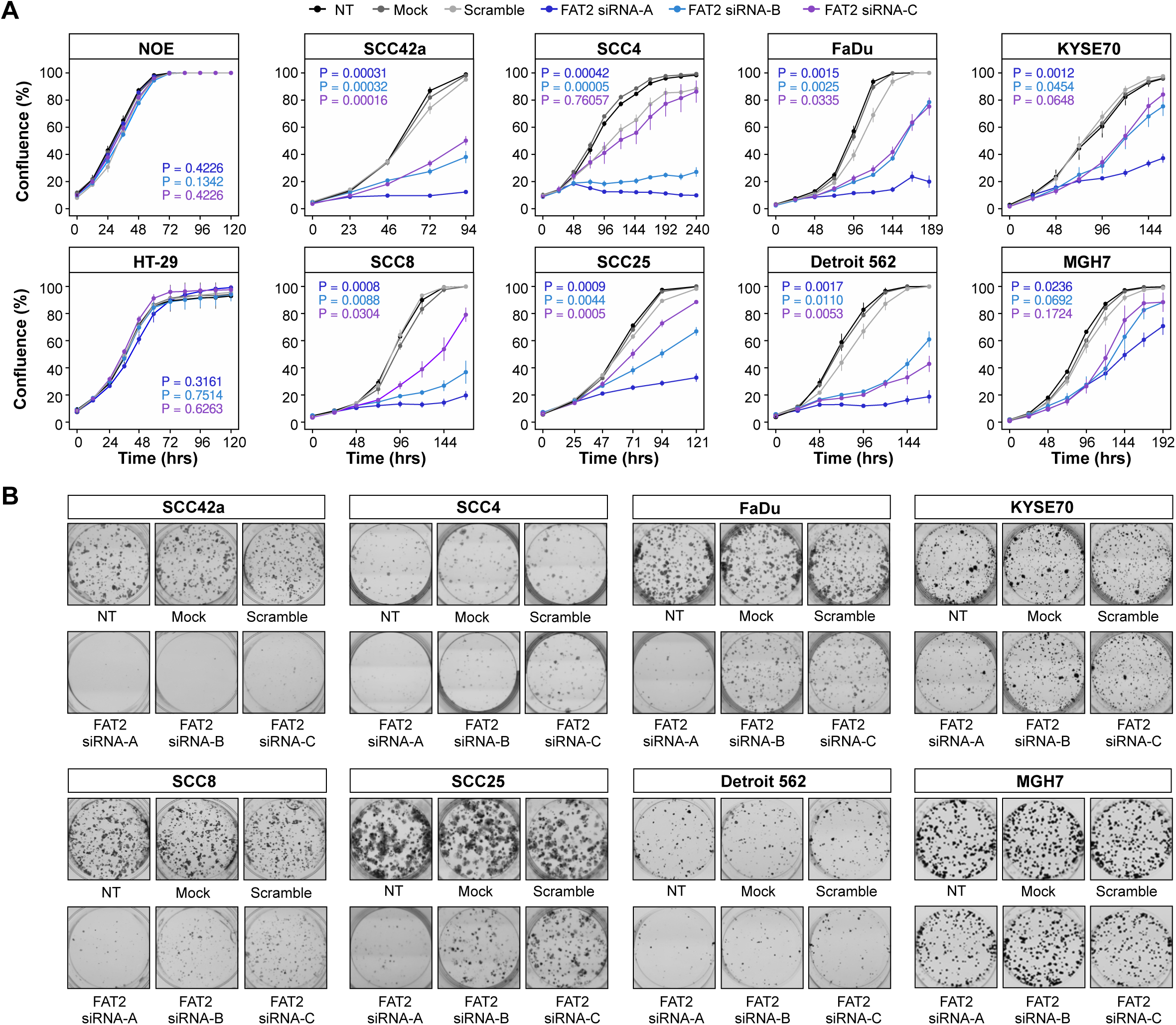
FAT2 is essential for HNSC growth and survival *in vitro*. **(A)** Proliferation of HNSC (SCC42a, SCC8, SCC25, SCC4, FaDU and Detroit 562), additional SCC (KYSE 70 and MGH7) and FAT2 negative (NOE and HT-29) cell lines following transient transfection with FAT2 targeting siRNAs (n = 3). NT = non-treated cells. Data are represented as mean ± SD and P-values were determined with an unpaired Student’s t-test against scramble at endpoint. **(B)** Colony formation assays of SCC cell lines following transient transfection with FAT2 targeting siRNAs. All data are representative of three independent experiments. See also **Figure S5**.

To obtain a stable system for further phenotypic evaluation, we generated a doxycycline (dox)-inducible shRNA expression system in two HNSC cell lines (SCC42a and SCC8). Incubation with low dose dox (1µg/mL) resulted in a rapid and reproducible knockdown of FAT2 protein, using two independent hairpins (**Figure 5A**). Akin to transient downregulation, shRNA-mediated knockdown of FAT2 had deleterious effects on *in vitro* HNSC growth and survival compared to control conditions (scramble) (**Figures 5B**, **5C**). To evaluate the *in vivo* consequences of FAT2 depletion, we subcutaneously engrafted SCC42a and SCC8 cells expressing dox-inducible shRNA into immunocompromised mice. Once tumors were established, FAT2 downregulation was initiated with the addition of dox in the drinking water, leading to a dramatic reduction of tumor size in both HNSC models compared to scramble controls (**Figures 5D, 5E**). Importantly, this FAT2 knockdown-mediated decline in tumor burden correlated with a statistically significant improved overall survival of mice engrafted with SCC42a HNSC cells (**Figure 5F**). Time-course IHC and immunoblotting analyses of harvested xenograft tumors following FAT2 depletion revealed substantial reduction in proliferation (Ki-67) and induction of apoptosis (TUNEL, cleaved PARP, cleaved caspase 3), but no measurable changes in tumor vascularization (CD31) compared to controls (**Figures 5G-5I**). Altogether, these data demonstrate that FAT2 is vital for HNSC xenograft tumor growth.

**Figure 5.**
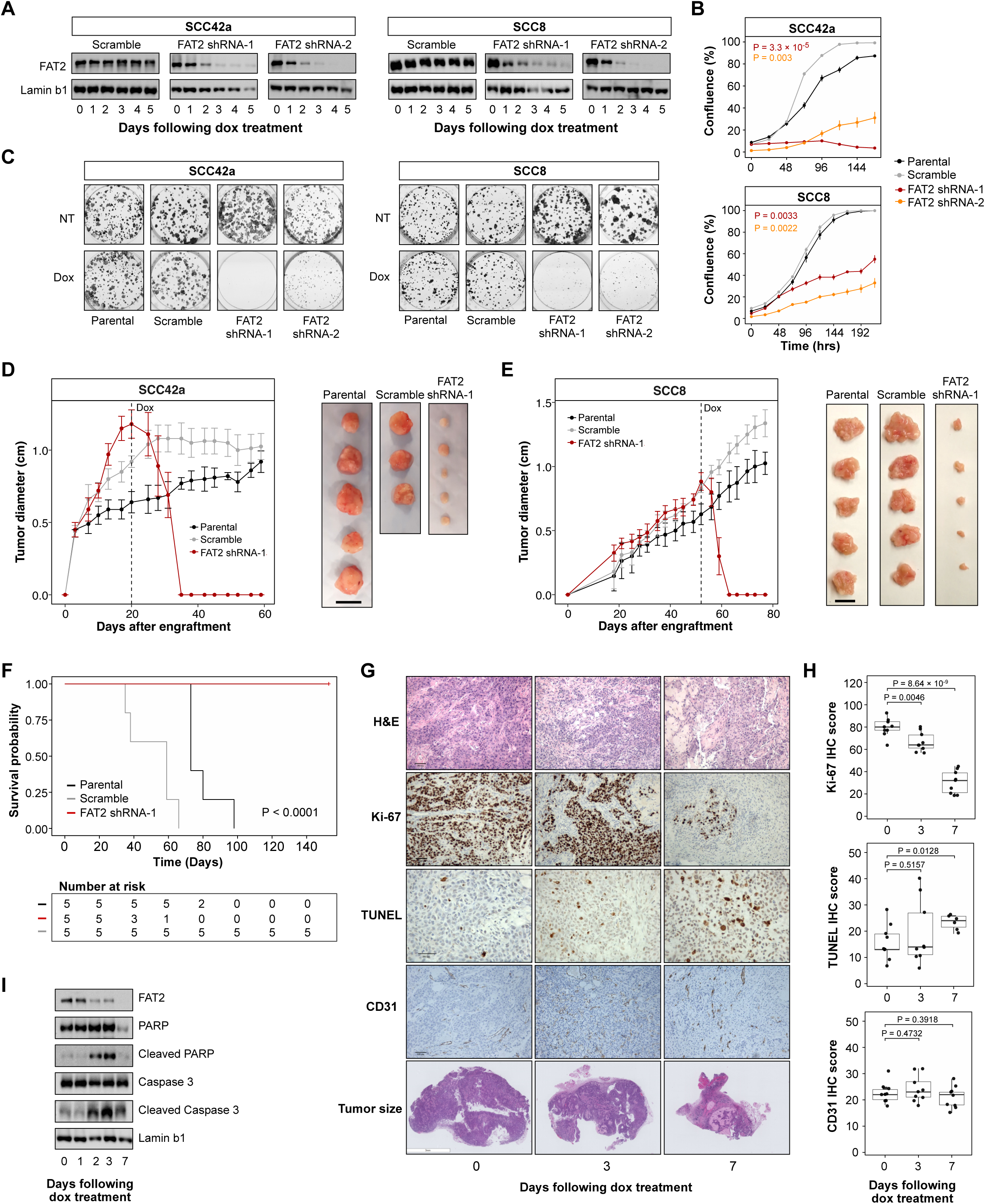
shRNA mediated depletion of FAT2 impairs *in vivo* HNSC growth. **(A)** Immunoblots of FAT2 abundance in a doxycycline (dox) inducible shRNA expression system established in HNSC cell lines SCC42a and SCC8. **(B)** Cell proliferation (mean ± SD) using the inducible shRNA expression system in SCC42a and SCC8 cells (n = 3). P-values were determined with an unpaired Student’s t-test against scramble at endpoint **(C)** Colony formation assays using the inducible shRNA expression model with the SCC42a and SCC8 lines. Representative of three individual experiments **(D & E)** *In vivo* growth curves (mean ± SD, left) and images of tumors harvested at endpoint (right) for subcutaneous **(D)** SCC42a and **(E)** SCC8 tumors with dox inducible shRNAs (n=5 for SCC42a parental and FAT2 shRNA 1, n=3 SCC42a scramble, n=5 for SCC8 parental, scramble and FAT2 shRNA 1). Dashed line indicates beginning of dox administration. Scale bar = 1 cm. **(F)** Kaplan-Meier survival curves of mice bearing SCC42a xenografts (n = 5). Statistical significance was calculated with a log-rank test. **(G)** Representative images of H&E and IHC evaluation of SCC42a xenograft tissues at defined time points after FAT2 shRNA induction. Scale bar: 100µm for H&E, Ki-67 and CD131, 200µm for TUNEL, 3mm for tumor size. **(H)** Images were scored for proliferation (Ki-67), apoptosis (TUNEL), and vascular density (CD31) (n = 9). P-values were calculated using an unpaired Student’s t-test against the values of day zero. **(I)** Immunoblots of cleaved PARP and Caspase 3 in FAT2 shRNA-induced tumors. Representative of three independent experiments.

### FAT2 mediates HNSC surface organization, adhesion and survival through integrin-PI3K signaling

Given the established role of cadherins in cellular adhesion and extracellular interactions^42^, as well as the large extracellular domain of FAT2 (over 440KDa), we hypothesized that the observed phenotypes are mediated by cell surface interactions. Consequently, we mapped the HNSC surfaceome as a function of FAT2 protein downregulation by performing CSC on the described dox-inducible shRNA SCC42a models three days post dox administration (**Figure 6A**) – a time point at which FAT2 is consistently depleted (**Figure S6A**) but just prior to detrimental phenotypes occurring in SCC42a cells. Major changes in plasma membrane protein composition were observed following FAT2 knockdown, with 79 proteins upregulated and 108 proteins downregulated, including FAT2 itself (**Figure 6B, Table S4**). Pathway analysis of the downregulated proteins revealed associations to cell adhesion, ECM-receptor interaction and PI3K-Akt signaling processes (**Figure 6C, Table S4**). Of particular interest, cell surface abundance of most detected integrins (9/12) decreased as a function of FAT2 protein depletion (**Figure 6D**). To expand on the observation of dysregulated adhesion related proteins, we measured adhesion of SCC42a cells to defined ECM components following FAT2 depletion. Knockdown of FAT2 resulted in reduced adhesion to collagen I (**Figures 6E**) and laminin (**Figures S6B, S6C**), with adherent FAT2-knockdown cells also occupying smaller surface areas compared to a Scramble controls (**Figures S6D, S6E**). In summary, our results suggest that reduction of FAT2 in HNSC impairs cellular adhesion via remodeling of the cellular surfaceome.

**Figure 6:**
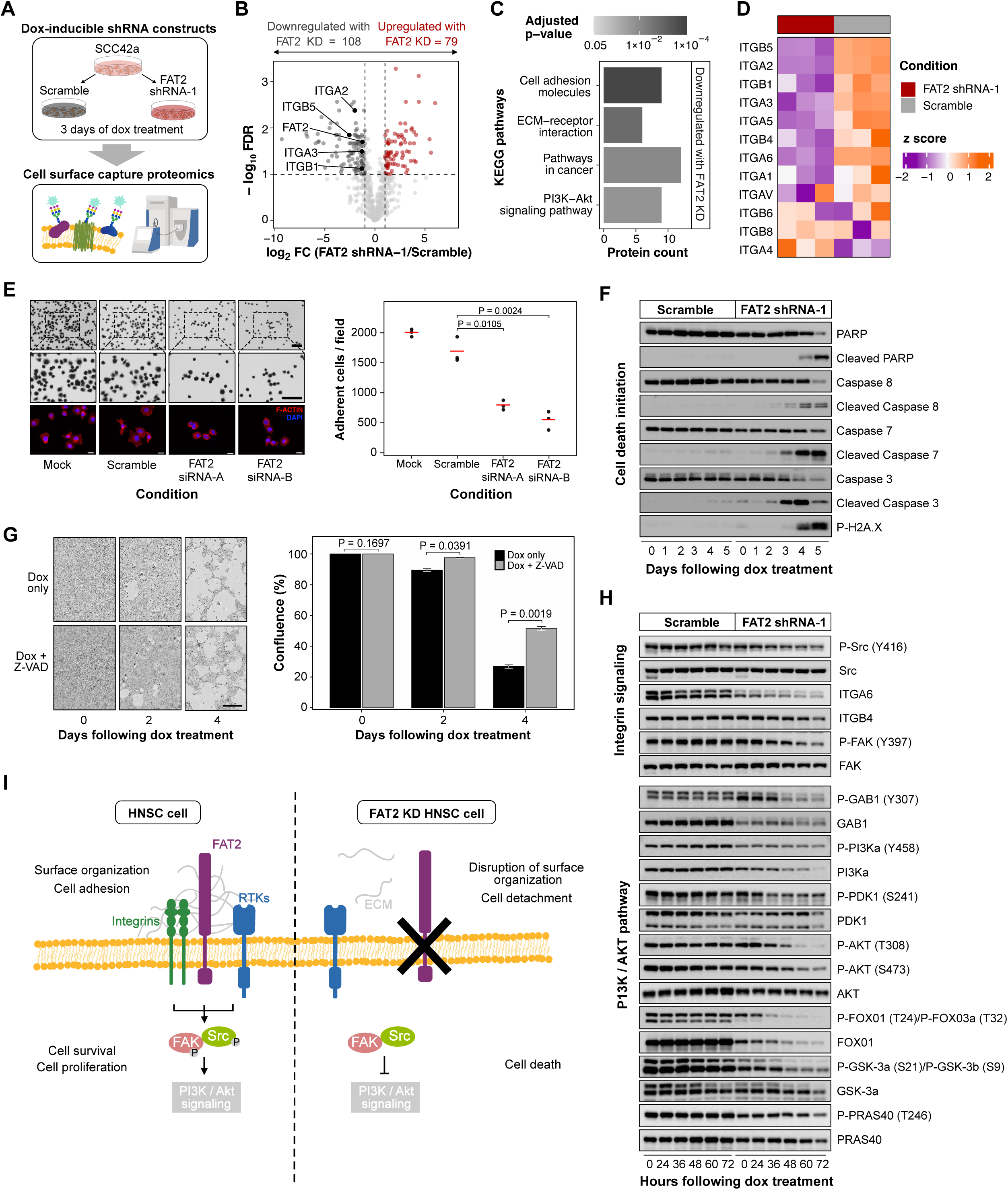
FAT2 mediates HNSC surface organization, adhesion and survival through integrin PI3K signaling. **(A)** Overview of cell surface proteomics performed on SCC42a cells following FAT2 depletion. Three replicates per condition. **(B)** Differential surfaceome expression following FAT2 depletion in SCC42a cells. Significantly changed proteins are indicated in black (downregulated) or red (upregulated). P-values calculated by unpaired Student’s t-test with FDR. **(C)** Select KEGG pathways significantly downregulated following FAT2 depletion. **(D)** Heatmap depicting quantitative changes in all detected integrins as a function of FAT2 depletion (z-scores). **(E)** Cell adhesion of SCC42a cells to collagen I following depletion of FAT2. Adherent cells are shown in the top and middle panels (Scale bar: 100 µm) and F-actin staining in red is shown on the bottom (Scale bar: 30 µm). Mean (red line) and individual data points (black) are visualized on the right (n = 3). P-values were calculated with an unpaired Student’s t-test and data are representative of three independent experiments. **(F)** Immunoblot analysis of apoptosis activation in response to FAT2 downregulation. Images are representative of three independent experiments. **(G)** Cell detachment of confluent SCC42a cells following FAT2 downregulation in the presence of the pan-caspase inhibitor Z-VAD. Representative images of cellular confluency (left) and confluency quantification (right) from three experiments. Scale bar: 400µm **(H)** Immunoblot analysis of the effects of FAT2 downregulation on integrin signaling and the PI3K/AKT signaling pathways. **(I)** Schematic of FAT2 driven surfaceome organization and resulting activation of cell survival and proliferation pathways. See also **Figure S6**.

We next aimed to elucidate the molecular signaling underlying FAT2 associated phenotypes in HNSC. Corroborating and extending our xenograft data (**Figures 5G-5I**), dox-mediated FAT2 reduction in SCC42a cells led to time-dependent activation (*i.e.* cleavage) of PARP, caspase-3, −7, and −8 and phosphorylation of histone H2A.X – hallmarks of cell death programs (**Figure 6F**). Consistently, treatment of high-density SCC42a cultures with a pan-caspase inhibitor, Z-VAD, decreased cellular detachment following FAT2 knockdown (**Figure 6G**), raising the possibility that FAT2-depleted cells die from anoikis – a detachment-dependent cell death pathway^43,44^. Since our cell surface proteomics analysis following FAT2 protein knockdown revealed noteworthy reduction of multiple integrins (**Figures 6B and 6D**) and dysregulation of the pro-survival PI3K-Akt pathway (**Figure 6C**), we used immunoblotting for detailed interrogation of these signaling cascades (**Figure 6H**). Knockdown of FAT2 resulted in a time-dependent decrease in protein abundance for major integrins (ITGA6, ITGB4) and reduced activation-dependent phosphorylation of Src kinase (Y416) and focal adhesion kinase (FAK; Y397), without reduction of total Src or FAK protein levels (**Figure 6H**, upper panel). Moreover, activation-specific phosphorylation of multiple components of the PI3K-Akt signaling pathway, which acts downstream of integrin signaling, was coordinately reduced upon time-dependent FAT2 protein depletion (**Figure 6H**, lower panel). Together, our data support a model whereby FAT2 protein is upregulated in HNSC and involved in cell surface protein organization and cell adhesion. Ultimately, this facilitates sustained activation of integrin-PI3K-Akt signaling pathways promoting cell proliferation and survival. Deletion of FAT2 in HNSC remodels the surfaceome to downregulate adhesion molecules resulting in cell detachment, impaired cell growth signaling and activation of cell death pathways (**Figure 6I**).

### FAT2 is a novel immunotherapy target for HNSC

Having validated FAT2 as a surface protein highly abundant in HNSC cells with limited expression in normal tissue and demonstrating its essentiality for HNSC growth, we aimed to evaluate proof-of-concept applicability of FAT2 targeted therapies. To this end, we repurposed our A088 FAT2 monoclonal antibody into a second-generation CAR backbone. The construct consisted of a CD8α signal peptide, the anti-FAT2 single chain variable fragment (scFv), a G4S linker, CD8α hinge domain, CD28 transmembrane and co-stimulatory domains and an intracellular CD3ζ signaling domain (**Figure 7A**). The CAR construct was separated from a truncated human EGFR tag by a P2A peptide. Normal donor-derived T cells were then transduced with lentivirus containing CAR constructs, assessed for transduction efficiency **(Figures 7B and 7C)** and co-cultured with FAT2 expressing HNSC cells (SCC42a or SCC8). Co-culture of FAT2 CAR T cells with HNSC cell lines at various effector/target ratios induced target cell cytotoxicity in a dose-dependent manner compared to co-culture with untransduced (UTD) T cells (**Figures 7D and 7E**). Co-culture of FAT2 CAR T cells with HNSC cells also led to an elevated production of cytokines (interferon [IFN]-γ and tumor necrosis factor [TNF]-α) (**Figures 7F and 7G**) and upregulation of T cell activation surface markers (CD25 and CD69) (**Figures 7H and 7I**) compared to UTD cells in both HNSC models. Together, these data reveal that FAT2 CAR T cells induce potent toxicity in FAT2 expressing HNSC cells and support the value of FAT2 as a novel immunotherapy target.

**Figure 7:**
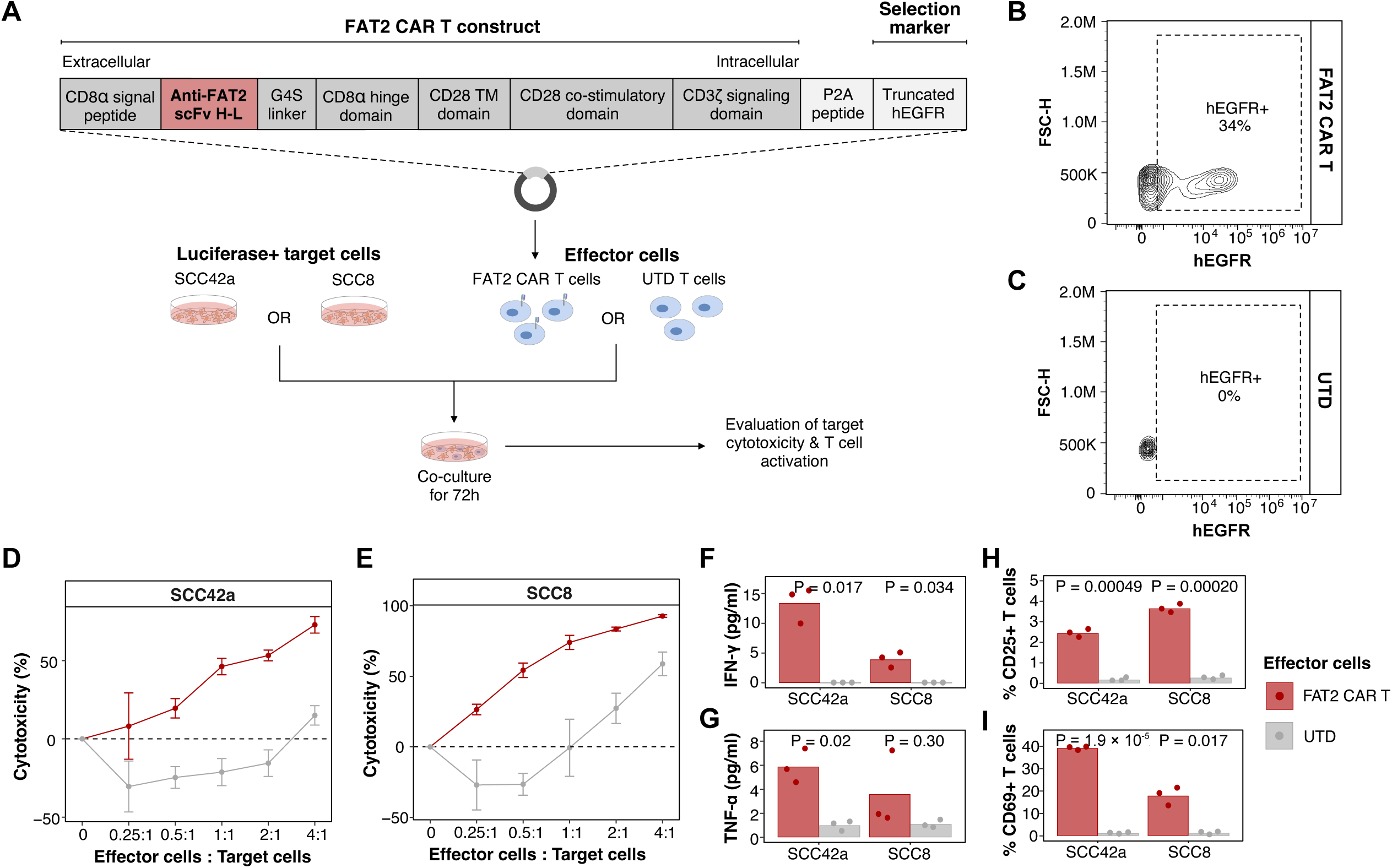
FAT2 is a novel immunotherapy target for HNSC. **(A)** Schematic of FAT2 chimeric antigen receptor (CAR) T construct and experimental design for co-culture experiments. UTD = untransduced negative control T cells. (**B & C**) Representative flow cytometry contour plots indicating hEGFR expression in **(B)** FAT2 CAR T cells and **(C)** UTD T cells to evaluate transduction efficiency. **(D & E)** Percent cytotoxicity of **(D)** FAT2 positive SCC42a and **(E)** SCC8 target cells co-cultured with FAT2 CAR T or UTD T cells at various effector:target ratios. Means and standard deviations from n=3 are plotted. **(F & G)** ELISA based quantification of T cell activation cytokines **(F)** interferon (IFN)-γ and **(G)** tumor necrosis factor (TNF)-α in supernatant following co-culture of head and neck squamous carcinoma (HNSC) cells with FAT2 CAR T or UTD T cells. **(H & I)** Flow cytometry analysis of **(H)** CD25 and **(I)** CD69 expressing T cells following co-culture with HNSC cells. **(F-I)** Means and individual data points are visualized (n=3). P-values from unpaired Student’s t-test.

## DISCUSSION

Despite their relevance as attractive drug targets, cell surface proteins are challenging to profile with common approaches (*e.g*. transcriptomics and global proteomics) and remain understudied. While we and others have extensively demonstrated the advantages of chemoproteomic enrichment of *N*-glycoproteins for cell surface profiling, most *N*-glycoproteomic surface discovery studies to date have been conducted in immortalized cancer cell lines or patient tissues^15,16,45–56^. Cell lines are advantageous for experimental manipulation and the considerable sample requirement traditionally required for *N*-glycoproteomic enrichment. Nevertheless, cell lines are susceptible to genetic drift and in some notable cases, poorly approximate patient tumors^57–59^. Direct investigations of patient tumors alleviates some of the challenges associated with cell line work, yet bulk tumor profiling is invariably prone to stromal contamination confounding molecular signals emanating from cancer cells^60^. In contrast to existing strategies, the unique target discovery platform described in this study is anchored in the use of PDX models and leverages the under-utilized power of PDX profiling to distinguish between cancer and the tumor microenvironment, thus directly addressing the limitations of profiling cell lines or patient tumors alone. Our pan-cancer analyses of 85 PDX *N*-glycoproteomes enabled the large-scale identification of cancer-derived surface proteins *in vivo*. We subsequently devised a PDX driven, multi-omic target discovery pipeline to prioritize 290 cancer enriched cell surface proteins displaying restricted expression in normal tissue as putative immunotherapy targets.

Our *N*-glycoproteomic PDX pipeline signifies the first step in target discovery to determine surface proteins with favorable expression profiles as potential candidates. As proof of principle for viable target identification, we sought to characterize and validate a novel PDX target candidate for HNSC. Considering that independent validation experiments (*e.g*. antibody generation, functional characterization *etc.*) are resource-intensive, we used *N*-glycoproteomics of HNSC patient tumors and cell lines to further prioritize high confidence PDX target candidates based on patient expression and cell surface localization, respectively. Despite the limitations of profiling only patient tumors or cell lines, we illustrate how such datasets can be used synergistically with the PDX *N*-glycoproteome atlas to provide additional confidence to PDX target candidates and selected protocadherin FAT2 as a strong candidate for further study. Evaluation of publicly available data and independent validation with a newly developed anti-FAT2 antibody in patient cohorts corroborated FAT2 enrichment in HNSC and its limited expression in normal tissue.

The Fat family of cadherins are adhesion molecules structurally conserved from Drosophila to vertebrates. In Drosophila, Fat proteins have been linked to planar cell polarity, collective cell migration and tissue rotation^61–64^. In vertebrates, the Fat family has expanded from two to four genes (*FAT1*, *FAT2*, *FAT3* and *FAT4*) and phylogenetic analyses suggest that mammalian *FAT2* is the most distinct and least studied family member^33,65^. The human *FAT2* gene was initially described in 1998^66^ and its protein product has been reported to localize at the cell-cell boundaries of keratinocytes^67^. Limited functional investigations of FAT2 implicate it in skin SCC cell migration^68^ and breast cancer invasion as part of the p63-induced transcriptional program^38^. The paucity of data regarding FAT2 further highlights the ability of our surface target pipeline to uncover under-appreciated cancer enriched surface proteins.

The extensive functional characterization presented here reveals that FAT2 is essential for *in vitro* and *in vivo* HNSC tumor growth. FAT2 knockdown leads to largescale remodeling of the surfaceome with widespread downregulation of a spectrum of adhesion molecules, including many integrins. Our work suggests that FAT2 depletion leads to impaired cellular attachment and downregulation of integrin-PI3K-Akt signaling, resulting in activation of cell death pathways, thus limiting HNSC growth. Considering the association of high FAT2 abundance with worse outcomes in HNSC, our data indicate that elevated FAT2 in HNSC and the associated cell surfaceome changes are part of a pro-survival program responsible for increased disease burden and adverse outcomes. A recently described model based on the amplification of SCC specific transcription factor networks via p63 and SOX2^69^ and previous identification of FAT2 as a transcriptional target of p63^38^ support such a view. Overall, our study sheds new light towards our understanding of FAT2 function in HNSC and alludes towards its potential broader role in SCC biology.

Towards clinical implementation of our findings, we approached FAT2 as a potential HNSC therapeutic target. By repurposing our newly developed monoclonal FAT2 antibody into a CAR T construct, we engineered FAT2 CAR T cells with potent cytotoxicity against two HNSC cell models. This proof-of-concept validation of FAT2 as a therapeutic candidate should fuel future studies on evaluating additional immunotherapeutic modalities (*e.g*. ADC or bispecifics) for FAT2 to assess their efficacy as therapeutic leads for HNSC and other SCCs. More broadly, our identification and verification of FAT2 as an actionable protein fortifies the capacity of our PDX *N*-glycoproteomic platform for therapeutic target discovery.

In summary, we established Glyco PDXplorer – a comprehensive resource of cancer-derived surface proteins detected *in vivo* and present a PDX driven, multi-omic discovery pipeline to prioritize putative therapeutic target candidates for multiple cancers. Our approach demonstrates potential for the development of antibody-based immunotherapies that go beyond the limited number of targets that are currently under investigation and improve cancer treatment with reduced adverse effects in patients. The data generated in this work is accessible via an interactive data web portal (http://kislingerlab.uhnres.utoronto.ca/glycoPDXplorer/) for others to leverage into new discoveries.

## ACKNOWLEDGMENTS

M.G. is supported by the Ontario Graduate Scholarship and Ontario Student Opportunity Trust Fund Awards. This work was funded by grants from the Canadian Institutes of Health Research (PJT 173487), the Canadian Cancer Society (705715) and the Canada Research Chair Program to T.K. We thank the Princess Margaret Living Biobank at University Health Network for generating LUSC, LUAD, COAD and PAAD PDX models. The Genotype-Tissue Expression (GTEx) Project was supported by the Common Fund of the Office of the Director of the National Institutes of Health, and by NCI, NHGRI, NHLBI, NIDA, NIMH, and NINDS. The results here are in whole or part based upon data generated by the TCGA Research Network.

## AUTHOR CONTRIBUTIONS

M.G., S.M-G., S.C.C, S.K., W.S., M.W., S.P., L.S., N.T., C.V., P.M., M.Topley, S.G., D.M., M.S., N.P., A.C., H.K., C.K., J.M., P.B., B.S., P.C., L.P., J.C., and A.R. designed experiments, performed experiments and contributed to data acquisition. M.G., S.M-G, V.I., A.K., L.Y.L., and T.K. analyzed data. M.G., S.M-G., and V.I. generated figures. T.K., S.K.S., I.S., E.C., F.L., V.S., L.A., D.G., W.X., S.S., R.K., and M.Tsao. supervised the study. M.G., S.M-G., and T.K. wrote the first full draft of the manuscript and all authors contributed to editing and approved the final manuscript.

## SUPPLEMENTARY INFORMATION

**Document S1:** Figures S1-S6, Note S1

**Table S1:** Clinical information & processed proteomics data pertaining to PDX *N*-glycoproteomics

**Table S2:** Normal tissue data metrics used for target prioritization

**Table S3:** Clinical information & processed proteomics data for HNSC tumor *N*-glycoproteomics

**Table S4:** Processed proteomics data for HNSC CSC experiments

**Table S5:** Patient characteristics and IHC data for TMA experiments

**Table S6:** LC-MS/MS method parameters

## STAR METHODS

### RESOURCE AVAILABILITY

#### Lead contact

Further information and requests for resources and reagents should be directed to and will be fulfilled by the lead contact, Thomas Kislinger.

#### Materials availability

All stable and unique reagents generated in this study are available from the lead contact upon request.

#### Data and code availability

- All mass spectrometry raw files acquired in this study are publicly available from UCSD’s MassIVE database (ftp://massive.ucsd.edu) under the dataset identifier #MSV000096571 and FTP link ftp://MSV000096571@massive.ucsd.edu. Processed proteomics data are available in this paper’s **Tables S1, S3 & S4**.
- A web application (http://kislingerlab.uhnres.utoronto.ca/glycoPDXplorer/) has also been created for browsing data.
- This paper does not report original code.
- Any additional information required to reanalyze the data reported in this paper is available from the lead contact upon request.

### ADDITIONAL RESOURCES

An accompanying web portal, available at http://kislingerlab.uhnres.utoronto.ca/glycoPDXplorer/, has been made freely accessible for researchers to explore and visualize *N*-glycoproteomics data from patient-derived xenografts. The platform was built with Python’s Django framework and integrates cutting-edge technologies like Bootstrap and D3.js for a seamless user experience while interrogating the Glyco PDXplorer data.

### EXPERIMENTAL MODEL AND STUDY PARTICIPANT DETAILS

#### Mouse strains and housing

Young adult (6-8-weeks old) immunodeficient NOD.Cg-Prkd^cscid^ Il2rg^tm1Wjl^/SzJ (NSG, Strain # 005557) mice from the Jackson Laboratory were used for experiments. Mice were housed in a modified barrier, specific pathogen-free facility in sealed negative ventilation cages (Allentown) in groups of maximum five mice per cage, at 22°C–24°C and a 12-h light/12-h dark cycle with food and water *ad libitum*. Animal experiments were performed according to guidelines from the Canadian Council for Animal Care and under protocols approved by the Animal Care Committee of the Princess Margaret Cancer Centre (Protocol numbers 5555, 1542 or 6396) or according to approved protocols from the Central Animal Facility at McMaster University (19-01-01). Clinical information for PDX models is indicated in **Table S1**.

#### Human tissues

##### HNSC N-glycoproteomics

As previously described^30^, HNSC tumor samples were collected from 87 patients with informed consent undergoing surgery at Toronto General Hospital, University Health Network. All experiments were approved by the Research Ethics Board at University Health Network (REB#18-5005). Patient characteristics are listed in **Table S3**.

##### HNSC IHC

Analysis of FAT2 expression in HNSC tumor tissue was conducted at the Princess Margaret Cancer Centre (PMCC, Toronto, ON, Canada) as a retrospective study in a cohort of two custom-designed Tissue Micro Arrays (TMAs) consisting originally of 117 HNSC patient samples obtained between 2003 and 2011. Studies were approved by the Research Ethics Board at University Health Network (REB#10-0725). Patient characteristics are indicated in **Table S5**.

#### Cell lines

The human laryngeal Squamous Cell Carcinoma (SCC) cell lines UTSCC8 and UTSCC42a were kind gifts from R. Grénman (Turku University Hospital, Turku, Finland). FaDu (pharynx SCC), Detroit-562 (pharynx SCC), SCC4 (tongue SCC), SCC25 (tongue SCC), and HT-29 (colorectal adenocarcinoma) cells were from ATCC. KYSE-70 (esophageal squamous carcinoma) and MGH7 (lung squamous cell carcinoma) were kind gifts from M. Tsao (Princess Margaret Cancer Centre, Toronto, Canada). Normal Oral Epithelium (NOE) cells were from Celprogen (Torrance, CA, United States). HNSC cell lines were grown in IMDM, KYSE-70 and MGH7 cell lines were grown in RPMI-60, and HT-29 cells were grown in McCoy’s 5A media. All media were supplemented with 10% FBS and PSG (20 U/ml penicillin, 20 U/ml streptomycin, and 60 µg/ml glutamine). NOE cells were cultured in NOE specific media according to the supplier’s indications. All cell lines tested negative for mycoplasma contamination.

## METHOD DETAILS

### Generation of patient-derived xenografts

#### COAD, LUAD, LUSC, PAAD models

Established PDX tumor models (COAD, LUSC, LUAD, and PAAD) were revived from cryopreservation and grown in NSG mice at the flank subcutaneous pocket, in accordance to previously described methods^70,71^. Once tumor sizes reached an average of 1 cm cubic, including skin fold, mice were anesthetized by inhaling isoflurane. Tumors were resected, flash-frozen and stored at −80°C until sample processing. Additionally, quality assurance was performed to evaluate concordance of PDX with matched patient tumor for histology and DNA fingerprinting. Studies were approved by the Research Ethics Board at University Health Network (REB#17-5518).

#### HNSC models

PDX models were generated as previously mentioned^30,72^. Under sterile conditions, HNSC patient tumors were cut into small pieces, and individual fragments were subcutaneously implanted into the flank of NSG mice. Tumor growth was monitored, and mice were euthanized when tumors reached 1.5 cm in diameter. PDX tumors were resected, flash-frozen and stored at −80°C until sample processing.

#### GBM models

As previously described^73^, GBM patient-derived brain tumor initiating cells (BTIC)^74–76^ were implanted intracranially into the right frontal lobes of 8-12 weeks old immunocompromised NSG mice, which were bred at the McMaster University Central Animal Facility. Briefly, mice were anaesthetized using 2.5% gas anaesthesia, isoflurane. A 1.0 cm vertical midline incision was made on top of the skull using a 15-blade. A small burr hole was then made on the skull, 2-3 mm anterior to the coronal suture and 3 mm lateral to midline using a drill held perpendicular to the skull. 10 μL of GBM BTIC suspension in PBS was injected 5 mm deep into the burr hole made in the frontal lobe using a Hamilton syringe. The incision was closed using interrupted stitches and sutures were sealed with a tissue adhesive. The mice were kept until they reached endpoint. The endpoint criteria were established based on a 20% reduction in body weight, worsening physical appearance, measurable clinical signs, abnormal or unprovoked behavior, and diminished response to external stimuli. Following euthanization, the entire brain was harvested, immediately snap-frozen on dry ice and stored at −80°C until sample processing.

### *N*-glycocapture

#### PDX tumors

All *N*-glycoproteomic experiments were approved by the Research Ethics Board at University Health Network (REB#11-0022). Flash-frozen PDX tumors were crushed using a cryoPREP Manual Dry Pulverizer (Covaris) and resuspended in SP3 buffer (100 mM HEPES, 1% SDS, 0.5% Triton X-100, 0.5% Tween 20, 0.5% NP-40). Samples were boiled for 10 min at 95°C followed by repeated sonication cycles using an indirect ultrasonic sonicator (Hielscher VialTweeter). Protein concentration was determined using the BCA assay (Pierce) according to manufacturer’s recommendations and 1 mg of protein lysate was used for subsequent sample processing. Disulfide bonds were reduced with 5 mM dithiothreitol (DTT) at 60°C for 30 min followed by alkylation of free sulfhydryl groups with iodoacetamide in the dark for 30 min at room temperature (RT). 5 mM DTT was used to quench the alkylation reaction. A magnetic bead based SP3 protocol^77^ was used for proteolytic digestion. Briefly, Sera-Mag Speedbeads (Cytiva) were mixed with protein lysates (10:1 (w/w) ratio) and 100% ethanol was added to achieve a 70% ethanol concentration. The mixture was shaken for 5 min at 1,000 RPM at RT to induce binding of proteins to the beads. Supernatant was discarded and beads were washed twice with 80% ethanol. Proteins were digested overnight with a trypsin/Lys-C mix (Promega) at a 1:50 protein ratio in 100 mM ammonium bicarbonate (NH_4_HCO_3_) at 37°C. The digestion was quenched with 1X protease inhibitor cocktail (Roche) the next morning and peptides were lyophilized in a SpeedVac vacuum concentrator. Enrichment of *N*-glycopeptides was performed as follows: lyophilized peptides were resuspended in coupling buffer (100 mM sodium acetate, 150 mM sodium chloride, pH 5.5) and glycans were oxidized using 10 mM sodium metaperiodate (NaIO_4_) at RT for 30 min in the dark. The excess NaIO_4_ was quenched with 20mM sodium thiosulfate for 15 min with vortexing. Peptides were coupled to hydrazide beads (Thermo) overnight at RT with constant rotation. The supernatant containing non-glycosylated peptides was discarded and the beads bound with *N*-glycopeptides were rigorously washed with coupling buffer, 1.5M NaCl, water, methanol, 80% acetonitrile and 100 mM NH_4_HCO_3_ to remove non-specific peptides. *N*-glycopeptides were enzymatically de-glycosylated and eluted from the beads with 5 U of PNGase F (Roche) in 100 mM NH_4_HCO_3_ at 37°C overnight. The de-glycosylated peptides were subsequently desalted using C18 stage tips (3M™ Empore™) and lyophilized. Peptides were solubilized with 0.1% formic acid (FA) in mass spectrometry (MS)-grade water and peptide concentration was determined using a NanoDrop 2000 (Thermo) spectrophotometer.

#### HNSC tumors

*N*-glycocapture was performed as stated above with minor changes. Prior to sample lysis, OCT was removed from shavings of OCT-embedded surgically resected tumor samples as previously described^30^. Specifically, samples were washed twice with 70% ethanol, twice with deionized water and once with 50mM ammonium bicarbonate followed by 5 min of air-drying at room temperature and resuspension in SP3 lysis buffer. Additional modifications include: 250 µg of protein lysate was used as starting input, magnetic hydrazide beads (Chemicell) were using for *N*-glycopeptide enrichment and SP2^78^ was used for desalting of peptides.

#### Cell Surface Capture (CSC)

For HNSC surface protein characterization experiments, each cell line (NOE, KYSE-70, MGH7, Detroit-562, FaDu, SCC4, SCC8, SCC25, and SCC42a) was seeded in triplicate in 150 mm plates and CSC was performed when cells reached 90% confluency. For FAT2 knockdown CSC experiments, SCC42a clones of FAT2-shRNA 1 or scramble transduced cells were seeded in triplicate in 150 mm plates and allowed to adhere for 24 h before doxycycline (1 µg/ml) treatment. Cells were grown in the presence of doxycycline for three days before CSC was performed. All cell surface labelling steps were performed on ice unless otherwise indicated. Cells were washed twice with washing buffer (PBS, 0.1% FBS), twice with labeling buffer (PBS, pH 6.5, 0.1% FBS), and mildly oxidized with 1 mM NaIO_4_ in 20 mL of labeling buffer with gentle rocking in the dark at 4°C for 15 min. Residual NaIO_4_ was removed with three washes of labelling buffer and cells were subsequently labelled with 5 mM biocytin hydrazide (Biotium, Fremont, CA, USA) in labeling buffer at 4°C with gentle rocking for 1 h. Excess biocytin was removed with three washes of labelling buffer. Cells were scraped off the plates and lysed in 1 ml of SP3 buffer for 10 min at 95°C followed by repeated sonication cycles using an indirect ultrasonic sonicator (Hielscher VialTweeter). Protein concentration was determined using the BCA assay (Pierce) according to manufacturer’s recommendations and 2 mg of protein lysate was used for subsequent sample processing. Protein denaturing and SP3-based digest was performed as described above for *N*-glycocapture. Enrichment of cell surface exposed *N*-glycopeptides was performed as follows: Peptides were mixed with 100 µl of Streptavidin Sepharose™ High Performance beads (GE HealthCare) in 100 mM NH_4_HCO_3_, pH 8.0, and incubated for 1 h at RT. Non-specific bead-binders were removed by extensive washing with 1 ml each of 0.1x invitrosol; 5 M NaCl; 100 mM sodium carbonate (Na_2_CO_3_), pH 11; 80% isopropanol, and 100 mM NH_4_HCO_3_, pH 8.0. *N*-glycopeptides were enzymatically deglycosylated and eluted from the beads using 5 U of PNGase F (Roche) in 200 µl of 100 mM NH_4_HCO_3_ at 37 °C overnight. Formerly *N*-glycosylated peptides were desalted using C18-based solid phase extraction stage tips (3M™ Empore™), lyophilized in a SpeedVac vacuum concentrator, and solubilized with 0.1% FA in MS-grade water. Peptide concentration was determined using a NanoDrop 2000 (Thermo) spectrophotometer.

#### MS-based proteomics

LC-MS/MS data for the PDX samples were acquired on an Orbitrap Eclipse Tribrid mass spectrometer (Thermo Fisher Scientific) directly coupled to a Neo Vanquish liquid chromatography system (Thermo Fisher Scientific). LC-MS/MS data for the HNSC patient tumors and CSC samples were acquired on an Orbitrap Q-Exactive and Q-Exactive HF (Thermo Fisher Scientific), respectively, coupled to an EASY-nLC 1000 nano-flow liquid chromatography system (Thermo Fisher Scientific). Peptides were loaded on a two-column setup using a PepMap^TM^ Neo Trap cartridge and Acclaim^TM^ PepMap^TM^ 100 column as trap columns for PDX and HNSC samples respectively, and a 50 cm EasySpray^TM^ ES903 column (Thermo Fisher Scientific) for peptide separation. Data acquisition was performed in positive-ion, data-dependent mode and specific LC-MS/MS parameters are listed in **Table S6**.

#### MS data processing

##### PDX

Raw files for each cohort (including OV PDX data from Sinha *et al.,*^14^ were searched separately with MaxQuant^79^ (version 2.4.2) against a concatenated human and mouse canonical plus isoform protein sequence database (Released 2022-04; 67,858 sequences) obtained from UniProt. Searches were performed with a maximum of two missed cleavages, and carbamidomethylation of cysteine as a fixed modification. Oxidation of methionine and deamidation of asparagine to aspartic acid were specified as variable modifications. The fragment ion peptide tolerance was set to ±20 ppm, and the parent ion mass tolerance was set to ±10 ppm. Match between runs was enabled in the search. The false discovery rate for the target decoy approach was set at 1% at site, peptide and protein levels. The Asn-AspSites.txt file from MaxQuant was used for subsequent analysis. Peptides detected with an asparagine deamidation modification within the *N*-glycosylation sequon *N-*[!*P*]*-S/T/C* (*N*= asparagine; [!*P*] = any amino acid other than proline; *S/T/C* serine, threonine or cysteine at the 2+ site) and with a localization probability > 0.8 were considered *N*-glycopeptides and used in subsequent analyses.

Species refinement was performed prior to PDX *N*-glycoprotein quantification **(Figure S1D)**. The initial species assignment of peptides was determined based on MaxQuant peptide detection against the combined canonical-isoform protein FASTA from UniProt. Peptides were assigned as either human-unique (H), mouse-unique (M) or human-mouse conserved (HM). To reduce the likelihood of possible false assignment due to potential database inconsistencies, species assignment was further refined by mapping detected *N*-glycopeptides to peptide sequences in (1) the UniProt TrEMBL predicted protein FASTA and (2) a custom protein-FASTA generated from a strain specific mouse genome to include variant peptides due to single nucleotide polymorphisms (SNPs) and indels. Briefly, germline SNPs and indels were obtained from the Mouse Genomes Project with GRCm38 VCFs downloaded from the European Variation Archive (accession: PRJEB43298) for the NOD/ShiLtJ mouse. Additionally, mouse variation VCF (GCF_000001635.24) for GRCm38.p4 was obtained from the dbSNP archive (v150). Germline SNPs and indels were mapped using VEP (v102) to ensembl GRCm38 GTF (v102), focusing on only primary chromosomes and amino-acid-changing mutations. moPepGen (v.0.11.3)^80^ was then used to generate peptides with amino acid alterations from all mutations and mutation combinations. All variant peptides were generated with trypsin digestion of up to 2 missed cleavages and peptide lengths 7-25, and max_variants_per_node = 1 and additional_variants_per_misc = 0 and otherwise default parameters. During refinement, all H peptides that mapped to a M or HM peptide in the TrEMBL or custom variant database were reassigned as HM and vice versa. For protein quantification, HM peptides were excluded from analysis and only H or M peptides were summed for quantification of human-derived or mouse-derived proteins, respectively. Proteins only detected by HM peptides were deemed indistinguishable and were omitted from further analysis. After evaluation of replicate variability **(Figure S1C)**, technical and processing PDX replicates were merged based on medians. Log_2_ transformed, median normalized *N*-glycoprotein intensities were used in subsequent analyses.

##### HNSC tumors

Raw files were searched with MaxQuant^79^ (version 1.6.3.3) using a UniProt human protein sequence database (Released 2022–04; 42,375 sequences). Search parameters and data processing was performed as described for PDX samples with minor changes. No species refinement was necessary for human HNSC tumor samples and *N*-glycopeptide intensities were summed as is for *N*-glycoprotein quantification. Log_2_ transformed, median normalized *N*-glycoprotein intensities were used in downstream analyses.

##### HNSC CSC

Raw files were searched with MaxQuant^79^ (version 1.6.3.3) using a UniProt human protein sequence database (Released 2022–04; 42,375 sequences). Search parameters and data processing was performed as described for PDX samples with minor modifications. No species refinement was necessary for human CSC samples and *N*-glycopeptide intensities were summed as is for *N*-glycoprotein quantification. Datasets for each experiment (*i.e.* HNSC cell line characterization and FAT2 KD) were filtered for proteins detected in 2/3 replicates of at least one cell line or condition. Log_2_ transformed, median normalized *N*-glycoprotein intensities were used in following analyses.

#### *In silico* species overlap analysis

Canonical protein FASTA sequences for human and mouse were obtained from UniProt (20,420 and 17,212 sequences for human and mouse, respectively). An *in silico* trypsin digestion was performed with the following rules: cleavage at the C-terminus of Arg/Lys, peptide lengths of 7-40 amino acids and no missed cleavages. Tryptic peptides were considered *N*-glycopeptides based on UniProt annotation data and the presence of the canonical *N*-glycosylation sequon whereas all other remaining peptides were deemed as global tryptic peptides. The percentage of both *N*-glycopeptides and global tryptic peptides conserved between species was calculated and P-values were determined using the Fisher’s exact test.

#### Identification of human PDX tumor type enriched clusters

Missing values were first imputed with random numbers drawn from a lower normal distribution (width = 0.2, down-shift = 1.8)^81^ and an ANOVA was performed to identify human-derived proteins that were differentially expressed between tumor types. Consensus clustering using ConsensusClusterPlus (v.1.64.0)^82^ was applied on proteins with a FDR < 0.05 (K = 8, finalLinkage = ward.D2, distance = Euclidean) to determine tumor type enriched human PDX *N*-glycoprotein clusters. Clusters were annotated by manual evaluation. Pathway analysis of proteins in each cluster was performed using g:Profiler^83^ and GO: biological processes.

#### Patient tumor type specificity analysis

##### TCGA

RNA-seq expression data from GBM, HNSC, LUSC, LUAD, PAAD, COAD and OV cohorts were queried using FirebrowseR (v.1.1.35)^84^ for all human PDX *N*-glycoproteins. Log_2_ transformed RSEM quantification was used for analysis. A Student’s t-test was used to determine statistical significances in one tumor type vs. the aggregate of the six remaining tumor types followed by Benjamini Hochberg procedure for multiple hypothesis correction. Gene-products with a log_2_ fold change > 2 and adjusted P-value < 0.05 for a tumor type against the other six were deemed tumor type enriched. For example, if a protein had a log_2_ fold change > 2 and an adjusted P-value < 0.05 in OV vs the collective GBM, HNSC, LUSC, LUAD, PAAD and COAD tumors, it would be annotated as OV enriched. ORs and Fisher’s exact P-values were determined to evaluate the degree of over-representation of each set of TCGA tumor type enriched proteins in each respective PDX human *N*-glycoprotein cluster and FDR was applied across all comparisons.

##### CPTAC

Proteomics data from GBM, HNSC, LUSC, LUAD, PAAD, COAD and OV cohorts re-processed with the BCM pipeline for pan-cancer multi-omics data harmonization^18^ were downloaded from the CPTAC “Pan-Cancer analysis” page on Proteomic Data Commons (Proteome_BCM_GENCODE_v34_harmonized_v1). Tumor type specific proteins were determined as stated above for TCGA except only proteins detected in > 50% of samples in each comparison group were retained for analysis.

#### Curation of clinical trial ADC and CAR-T targets for solid tumors

##### ADC targets

A list of all ADC clinical trials as of April 1, 2023 was obtained from the supplementary table of Dumontet *et al.,*^25^. These trials were manually filtered for trials of phase 1 or greater and focused on treating solid tumors. ADCs in these remaining trials targeted at least one of 34 unique protein targets.

##### CAR T targets

A list of all phase 1 or higher clinical trials on ClinicalTrials.gov which matched the condition search terms of “carcinoma” OR “cancer” OR “glioblastoma” and the intervention search terms of “CAR-T” OR “chimeric” was downloaded on July 12, 2023. These trials were manually inspected to retain trials which indeed were for CAR T therapies and focused on treating solid tumors. The remaining 277 trials evaluated CAR T therapies targeting at least one of 58 unique protein targets.

#### Normal tissue toxicity scoring

Three categories of publicly available normal tissue data were sourced and integrated:

1) Datasets that sampled normal tissue from the entire human body (*i.e.* pan-tissue): Normalized MS-based proteomics data on GTEx was obtained from the supplementary table from Jiang *et al.,*^20^. Processed median gene-level TPM per tissue profiled by GTEx was downloaded from the online GTEx data portal (2017-06-05-V8)^19^. Normal tissue IHC data were downloaded from Human Protein Atlas (V20.1)^5^.
2) Datasets that evaluated different regions or cell types from a normal human brain: Normalized TPM and IHC detection from various normal brain cell types was downloaded from Human Protein Atlas (V23.0)^22^. MS-based proteomics data was obtained from the supplementary table from Tushaus *et al.,*^21^.
3) Datasets that investigated different regions or cell types from a normal human heart: Processed cardiomyocyte CSC data from different regions of the heart profiled by Berg Luecke *et al.,*^23^ were obtained from the corresponding author of the study. MaxQuant search results for MS-based data from Doll *et al*.,^24^ were downloaded from PRIDE (PXD006675) and filtered to remove contaminants and reverse peptide hits.

To enable evaluation of normal tissue expression across various types of data (*e.g.* LC-MS/MS, RNA-seq and IHC), data metrics were classified as “*qualitative”* if binary absence or presence of a gene-product in samples was evaluated (*e.g.* IHC positive detection) or *“quantitative”* if the abundance of a gene-product was measured (*e.g.* RNA TPM) (**Figure S2C**, **Table S2**). For qualitative data metrics, the frequency of sample detection was calculated for each protein. For quantitative data metrics, the median abundance across all tissue or cell types was determined for each protein. In the event where a dataset contained multiple replicates per tissue or cell type, replicates were merged based on medians. A protein received a score of 1 if it was detected in < 50% of samples in qualitative datasets or if its median abundance was below the median of all PDX human proteins in quantitative datasets. Scores were summed for each tissue type category (*i.e.* pan-tissue, brain or cardiac). As there were four data metrics per tissue type category, scores ranged from zero to four. A score of zero meant the protein had low abundance and/or low detection frequency in zero out of four data metrics and a score of four indicated that the protein had low abundance and low detection frequency in all four datasets for the respective tissue type category.

#### PDX target prioritization pipeline

Two stages comprise the target discovery pipeline **(Figure 2C)**. First, Glyco PDXplorer was leveraged to prioritize cancer-enriched surface proteins. For each tumor type, human PDX *N*-glycoproteins (*i.e.* cancer-derived) were filtered for proteins that had a predicted surface localization (Surface Prediction Consensus [SPC] score^85^ >0) and were highly abundant in PDXs of the respective tumor type (detected in ≥ 50% of samples and/or median intensity of the protein was higher than the median protein intensity of the entire PDX dataset for a specific tumor type). The next stage involved the integration of multi-omic published normal tissue data to select proteins with limited abundance across all normal tissues and specifically in essential organs (*i.e.* heart and brain) to minimize the likelihood of normal tissue toxicity. To this extent, proteins were required to have a pan-tissue score ≥ 2, cardiac score ≥ 2 and neuro score ≥ 2. Two hundred and ninety unique surface proteins were determined to be cancer-enriched with limited expression in normal tissue in at least one of seven tumor types and represent putative therapeutic targets (*i.e.* PDX target candidates).

#### HNSC target verification and ranking

We appended two additional *N*-glycoproteomic stages to our PDX target discovery pipeline for the verification of HNSC target candidates. First, candidates were required to be detected in >50% of HNSC tumors processed by *N*-glycocapture to validate reproducible detection of targets in patients. Second, CSC was used to provide experimental evidence of a cell surface localization in HNSC and thus, candidates were required to also be detected on the surface of >50% HNSC cell lines profiled. The remaining 31 target candidates represent highly abundant HNSC-derived surface proteins with reproducible detection in patients and limited expression in normal tissue. A rank-sum approach was used to order candidates based on the frequency of detection and abundance in both HNSC PDX tumors and patient tumors as well as the log_2_ fold change in surface abundance between each SCC cell line and NOE. The top ranked protein, FAT2 was selected for further evaluation.

#### Immunofluorescence

HNSC cell lines, SCC8 and SCC42a, were seeded in 24-well plates (10,000 cells/well) containing sterile glass coverslips at the bottom. Cells were grown overnight, washed twice with PBS and fixed with fresh 4% paraformaldehyde in PBS at 4°C for 10 min. After three 5 min washes with PBS, cells were permeabilized for 2 min with 0.1% Triton-X100 in PBS (PBSt), washed three times with PBS, and blocked with 1% BSA in PBS for 1 h at RT. Coverslips were incubated overnight with a 1:50 dilution of FAT2 antibody (Santa Cruz) in blocking media, washed three times with PBS and exposed for 1 h to a secondary goat anti-mouse antibody coupled to Alexa Fluor-488 (Abcam). Coverslips were washed three times with PBSt, stained with 1 µg/ml DAPI, and mounted for analysis using an FV1000 confocal microscope (Olympus).

#### Immunoblotting

Whole cell lysates were prepared using RIPA buffer (50 mM Tris-HCl pH 8, 150 mM NaCl, 5 mM EDTA, 1% NP-40, 0.1% SDS), supplemented with Pierce^TM^ Protease Inhibitor Tablets/Pierce^TM^ Phosphatase Inhibitor Mini Tablets (Thermo Fisher Sci.). Protein extracts (20 µg) for FAT2 detection were resolved on a 7% SDS-PAGE gel and electro-transferred overnight onto a PVDF membrane. All other proteins were resolved on 8-15% SDS-PAGE gels. Membranes were blocked for 1 h in 5% skim milk TBST (0.1% Tween 20 in TBS) and probed overnight (4°C) with primary antibodies (refer to **Key Resources Table**). Following standard Western blot protocols, membranes were subsequently exposed to appropriate HRP-conjugated secondary antibodies (Cell Signaling Technology) and immunoreactive bands visualized with SuperSignal West Femto Maximum Sensitivity Substrate (Pierce) with a MicroChemi imager (Bio-Imaging Systems).

#### Development of FAT2 monoclonal antibodies

Monoclonal anti-FAT2 antibodies were generated by phage display in collaboration with adMare BioInnovations (Vancouver, BC, Canada). Briefly, the full-length extracellular domain sequence of human FAT2 (NP_001438.1/ UniProt Q9NYQ8) was divided into 6 smaller truncates and cloned. Two of these truncates, CAD29-TM and CAD18-24, were successfully purified and subsequently cloned as Fc fusion proteins. Immunization of Balb/C mice was performed with Fc-fusion protein or a FAT2 DNA expression plasmid. Spleens were harvested from all animals from both immunization protocols after ELISA serum titration and used to build a phage display library using a c-Myc tag for detection and a HIS6 tag for purification. After three rounds of selection, 15 unique clones were identified as binding to CAD29-TM, and 1 binding clone to CAD-18-24. All 16 binders were selected for reformatting to IgGs. Generated IgGs were characterized by their binding (ELISA, flow cytometry), affinity (Octed Red96e, ForteBio), epitope competition (Binning), internalization, and binding to a panel of membrane proteins (Membrane Proteome Array screening). Afterwards, clones were selected for testing by Western blot, immunocytochemistry (SCC42a cells), and immunohistochemistry on SCC42a-derived xenograft tissues. Finally, the top antibody candidate, A088, was selected for testing of FAT2 detection by IHC on commercially available FFPE human cerebellum (cat# T2234039) and liver tissue (cat# T2234149), both from BioChain, and a human adult normal tissue FFPE TMA (NBP2-78113, Novus Biologicals corp.) with 62 different tissue cores, spanning 31 different tissue types. An anti-EGFR antibody (1:500, Cell Signaling Technology) and a mouse IgG2a isotype (10 µg/ml, Abcam), were used as positive and negative controls, respectively. For a more detailed information on FAT2 antibody development see **Note S1**.

#### Immunohistochemistry analysis of patient samples

Immunohistochemistry (IHC) for FAT2 detection in tumor specimens was performed on a HNSC TMA custom-designed cohort at Princess Margaret Cancer Centre containing both tumor cores and adjacent normal tissue, when available. Initially, IHC was performed with a commercially available monoclonal FAT2 antibody (Santa Cruz Biotechnology). Briefly, tissue sections were deparaffinized with xylene, rehydrated through ethanol, and microwave-antigen retrieved with citric buffer (10 mM Sodium Citrate, 0.05% Tween-20, pH 6.0). The TMA was subsequently blocked with 3% hydrogen peroxide, followed by 10% Normal Goat Serum (NGS) in TBST (Tris Buffer Saline, 0.1% Tween-20). Afterwards, sections were incubated overnight at 4°C with the mouse monoclonal anti-FAT2 antibody. The TMA was subsequently processed with the EnVision(TM) + Dual Link System HRP DAB+ kit (Dako). Since FAT2 was found to be expressed in both the cell membrane and the cytoplasm of interpretable tumors, evaluation of expression levels of FAT2 in the two cellular structures was performed separately. Levels of FAT2 expression were evaluated for subcellular localization, staining percentage (proportion), and a 4-tiered score system for intensity (0, 1+, 2+, and 3+). Immunostaining score was then calculated by multiplying the proportion and intensity scores. The combination of membranous and cytoplasmic staining was added as the final score. FAT2 staining was evaluated at least twice, and all scores were performed blinded to clinical information. A Student’s t-test was used to evaluate statistical differences in FAT2 IHC scores between HNSC tumor and normal samples. Additional validation of FAT2 expression in the TMA was performed with the custom mouse-formatted anti-human FAT2 antibody, A088, at a 10 µg/ml concentration on the same TMA cohort. IHC and stain evaluation was performed as described above for the Santa Cruz Biotechnology antibody. Up to 13 (11%) TMA samples were considered uninterpretable due to lack of unequivocal tumor cells or loss of tissue during technical procedures and were excluded from analyses.

#### Survival analysis of tumor IHCs

FAT2 IHC scores were median dichotomized to define FAT2 high and low IHC patients. A log-rank test was used to test statistical significance for overall survival (OS) and recurrence-free survival (RFS) and Kaplan-Meier curves were visualized using the packages survival (v.3.5.5) and survminer (v.0.4.9) in R (v.4.3.0). Univariate and multivariate analyses were performed by Cox proportional hazard models. A two-tailed Student’s t-test was used, with a P-value < 0.05 as a significant level and analyses were performed using SAS 9.4 (SAS Institute, Cary, NC).

#### Evaluation of FAT2 expression in normal tissue datasets

Normal tissue IHC data were downloaded from Human Protein Atlas (v.23.0)^5^. To aggregate data from multiple cell types into a tissue level analysis, the maximum IHC detection level (e.g. “High”, “Medium”, etc.) across all cell types stained for a particular tissue type was retained for analysis and visualization. To evaluate spatially resolved expression of FAT2 in normal human skin, raw LC-MS/MS data pertaining to human subcutis, dermis, inner epidermis (stratum basale) and outer epidermis (stratum corneum) samples^34^ were downloaded from PRIDE (PXD019909) and searched with MaxQuant (version 1.6.3.3) using a UniProt human protein sequence database (Released 2022–04; 42,375 sequences). Search parameters were similar as stated above except without deamidation of asparagine to aspartic acid specified as a variable modification. Median normalized iBAQ values were used for analysis. To investigate normal brain cell type expression of FAT2, processed scRNA-seq data and cell type “supercluster” annotations were downloaded from the GitHub repository associated with Siletti et al.,^35^ and the package loomR (v.0.2.1.9000) was used to process the data in R (v.4.3.0). Data were used as is to determine the percentage of cells in each cluster that expressed FAT2. Data were log_2_ transformed to visualize FAT2 expression across the cell type superclusters.

#### FAT2 transient downregulation

Downregulation of FAT2 was achieved through a modified reverse-transfection protocol recommended for Lipofectamine RNAiMAX (Invitrogen). Transfection conditions were optimized for each cell line using a TYE 563 fluorescent-labeled transfection control (Origene, SR30002). siRNA duplex Control scramble and FAT2 siRNAs (all from Origene) were diluted in OptiMEM and mixed with Lipofectamine RNAiMAX transfection reagent at half the manufacturer recommended concentration. The mixture was deposited at the bottom of multi-well cell culture plates such as to achieve a final concentration of 1 nM siRNA upon addition of a cell suspension. FAT2 siRNA sequences were:

FAT2 siRNA-A: 5’- CCAAUGCUCAGAUCACAUAUUCUCT -3’
FAT2 siRNA-B: 5’- GGAGAGUACAACAUCCUAACGAUCA -3’
FAT2 siRNA-C: 5’- GCACUACACCUGAGAGCAACAAGGA -3’
Scramble siRNA: 5’- CGUUAAUCGCGUAUAAUACGCGUAT -3’

Unless otherwise specified, all experiments included, in addition to the scramble control, non-treated (NT), and cells treated with transfection reagents but no siRNA (Mock) controls.

#### FAT2 inducible downregulation

Following the supplier’s protocol, control (scramble) and FAT2 shRNAs were cloned into the pLKO-Tet-On lentiviral vector^86^ (a gift from Dmitri Wiederschain, Addgene plasmid # 21915; http://n2t.net/addgene:21915; RRID:Addgene_21915), where shRNA expression is repressed by a constitutively-expressed TetR protein. Upon addition of dox to the growth media, shRNA expression is triggered resulting in target gene knock-down. shRNA sequences were:

FAT2 shRNA-1: 5’- TACTACTGGTTGACGGTATTA -3’
FAT2 shRNA-2: 5’- TGCTCAGATCACATATTCTCT -3’
Scramble shRNA: 5’- CAACAAGATGAAGAGCACCAA -3’

For virus production, HEK 293T cells were seeded in low antibiotic (0.1x Pen/Strep) DMEM media at a density of 1.0 × 10^6^ cells/well in 6-well plates. 24 h later, cells were co-transfected with 850 ng of packaging plasmid psPAX2 (psPAX2 was a gift from Didier Trono, Addgene plasmid # 12260; http://n2t.net/addgene:12260; RRID:Addgene_12260), 350 ng of envelope plasmid VSV-G^87^ (pVSV-G was a gift from Akitsu Hotta, Addgene plasmid # 138479; http://n2t.net/addgene:138479; RRID:Addgene_138479), and 850 ng of pLKO-Tet-On plasmid in OptiMEM using X-tremeGENE 9 DNA transfection reagent (Roche) according to the manufacturer’s instructions. The day after transfection, media were replaced with viral harvest media (1% BSA, DMEM) and cells were incubated for an additional 24 h. Lentiviral supernatants were collected, passed through a 0.45 µm filter, and used immediately or stored at 4°C for a few days. For viral transduction 60,000 target cells/well were seeded in 6-well plates containing 8 µg/ml polybrene (Sigma). On the same day, and after cell adherence was confirmed, 100 µl of viral suspension was added, and cells were incubated overnight. Transduced cells were selected by treatment with puromycin (2 µg/ml) in full growth media for 48h. Surviving cells were collected for single cell cloning and characterization for subsequent experiments.

#### Cell proliferation, colony formation, and cell detachment assays

All proliferation experiments were carried out in triplicate in 96-well plates, where 500-1,000 cells were added per well and cell growth monitored using an Incucyte S3 Live-Cell Analysis Instrument (Sartorius). For experiments performed with dox-inducible shRNA expression clones, dox at a 1 µg/ml concentration in fresh media was replenished daily. Analyzed results are reported as percentage of Phase Object Confluence (POC). A Student’s t-test was used to assess statistical differences in in POC measurements at endpoint between each FAT2 siRNA or shRNA against the respective scramble. Colony Formation Assays (CFA) were performed in triplicate in 6-well plates at a density of 500-700 cells/well. For transient FAT2 downregulation, plates were left undisturbed for 10-14 days after siRNA transfection. For FAT2 inducible downregulation, 500-700 transduced cells were seeded in triplicate and treated with 1µg/ml dox on the same day, after attachment was confirmed. Dox was replenished every 2-3 days. Afterwards, plates were fixed and stained with 0.01% crystal violet, 20% methanol in PBS at RT. Colonies were counted manually and a Student’s t-test was used to evaluate statistical differences in colony number between each FAT2 siRNA or shRNA against the respective scramble. For experiments performed at confluence, cells transduced with the FAT2 shRNA-1 were seeded in triplicate at a density of 60,000 cells per well in 96-well plates. Typically, cells reached full confluence 24 h after seeding. At that time dox (1 µg/ml) alone, or in combination with the pan caspase inhibitor Z-VAD-FMK (10 µM), was added and cells were monitored daily for the duration of the experiment. Media was changed every 24 h and plates were scanned in an Incucyte S3 immediately afterwards. Analyzed results are reported as percentage of POC. A Student’s t-test was performed to investigate statistical differences in POC measurements between cells treated with and without Z-VAD-FMK at various timepoints following doxycycline treatment.

#### FAT2 *in vivo* xenograft experiments

Xenografts were generated by injecting subcutaneously into the flank of each animal 1.0 × 10^6^ cells of interest resuspended in 100 µl of equal volume reduced growth factor Matrigel (Corning) and serum-free media. Groups of five animals were injected each with parental, scramble, or FAT2-shRNA1 transduced cells. Tumors were measured biweekly using calipers and the longest perpendicular measurements used to report tumor size. Dox treatment, administered in drinking water (1 mg/ml in 20% sucrose), was started when control xenografts reached a length of 0.5 cm. Experiments were terminated when animals of any group reached endpoint - defined as a tumor size of approximately 15 mm in length - as recommended by the animal care committee. At endpoint animals were sacrificed with CO_2_ and tumors were removed, photographed, measured, and weighed. Samples from each tumor were paraffin embedded or flash frozen for subsequent immunohistochemistry or immunoblot analyses.

#### Differential expression of FAT2 CSC KD data

Missing values were imputed with lower tail imputation (downshift 1.8 s.d and width of 0.5 s.d)^81^. Log_2_ fold changes were calculated by the difference in means between FAT2 shRNA-1 and scramble transduced SCC42a cells and a Student’s t-test followed by Benjamini-Hochberg procedure for multiple hypothesis testing was used to determine statistically significant differences. Proteins with a log_2_ fold change > |1| and an adjusted P-value < 0.1 were considered significantly dysregulated in response to FAT2 knockdown. Pathway analysis was performed on proteins significantly upregulated and downregulated in response to FAT2 knockdown using g:Profiler^83^ and KEGG annotations.

#### Cell Adhesion to Extracellular Matrix Components

Adhesion experiments were performed in triplicate in 24-well plates covered overnight either with Collagen I (5 µg/cm2 in 0.02 N Acetic acid) or Laminin (2 µg/cm2 in serum-free media, SFM). Plates were washed twice with PBS immediately before the experiment. SCC42a cells were reverse transfected with Scramble, FAT2 siRNA-A, or FAT2 siRNA-B, and 48 h later trypsinized, counted, and seeded at a density of 1 × 10^5^ cells/750 µl SFM/well and allowed to adhere 20 min for Collagen I, and 45 min for Laminin. Afterwards, wells were washed twice with PBS and fixed and stained with 0.01% crystal violet, 20% methanol in PBS at RT. Cell numbers and relative sizes were analyzed with aid of the FIJI software^88^. A Student’s t-test was used to investigate statistical differences in number of adhered cells and cell size between cells treated with each siRNA against scramble.

#### Generation of FAT2 CAR T cells

The sequence of the scFv from the FAT2 monoclonal antibody was cloned into a second-generation CAR backbone as previously described^89^. Briefly, gene blocks were ordered from Twist Biosciences and cloned downstream of the EF1alpha promoter. The CAR construct contained a CD8a signal peptide, FAT2 vH-vL, G4S linker, CD8a hinge domain, the CD28 transmembrane/co-stimulatory domain followed by the CD3zeta intracellular signaling domain and a P2A peptide separating the CAR construct from a truncated human EGFR sequence. Human peripheral blood mononuclear cells from healthy donors were thawed from cryostorage in ImmunoCult-XF medium (Stem Cell Technologies # 100-0956) containing 100 U ml^−1^ IL-2 in the presence of ImmunoCult™ CD3/CD28/CD2 T cell activator (Stem Cell Technologies # 10970) in 96 well plates overnight (150, 000 per well). The following day, T cells were transduced with lentivirus encoding the CAR construct at an MOI ∼1.0. The following day, media was topped up with ImmunoCult containing IL-2 and T cells expanded for 7 days before confirming expression of the CAR construct *via* staining for truncated EGFR. CAR T cells were then cryogenically stored the following day only to be revived and expanded for use in assays between days 12-14 post-transduction.

#### CAR T lentivirus production

2 × 10^7^ HEK293T cells were plated in a T-75 flask in DMEM + 10% FBS + 1x non-essential amino acids (D10). The following morning media was removed and replaced with fresh, pre-warmed D10 containing sodium butyrate (D10V) 1 h prior to transfection. Gag/ Pol (5.3 µg), Rev (2.6 µg) and VSVg (2.6 µg) were mixed with the FAT2 CAR vector (10.6 µg) and polyethyleneimine (PEI) in a ratio of 3:1 µg PEI: µg DNA. The following morning the media was removed and replaced with fresh D10V. This media was collected the following day and concentrated 300x by ultracentrifugation at 20, 000 rpm for 2 h at 4 °C. The supernatant was removed and the virus resuspended overnight in ImmunoCult-XF and then frozen at −80 °C.

#### Flow cytometry

##### Transduction efficiency

Expression of the CAR construct in transduced T cells was confirmed by staining for human EGFR (1:50). 250,000 T cells were collected from actively growing expanding cultures and stained for human EGFR for 15 min at room temperature. Cells were stained with 7AAD to ensure that only viable cells were being analyzed. 10,000 T cells were analyzed, and the transduction efficiency determined as the proportion of hEGFR positive cells. Data were analyzed with FlowJo^TM^ (v 10.0).

##### T cell activation

To assess T cell activation, 500, 000 target cells were cultured with T cells at an E:T ratio of 1:1 in triplicate for 72 hours in 50% target cell medium/50% ImmunoCult-XF medium. After 72 hours, T cells were removed from each well and cell culture supernatant saved for ELISAs below. T cells were washed 1x with PBS and stained for 15 minutes at room temperature in the dark with antibodies to CD3 (1:50), CD25 (1:50) and CD69 (1:20) and EGFR. T cells were then washed 1x with PBS and resuspended in 300 µL containing 7AAD to ensure only viable cells were analyzed. Data were analyzed with FlowJo^TM^ (v 10.0).

#### *In vitro* cytotoxicity assays

Luciferase expressing variants of SCC8 and SCC42a were generated using the lentiviral pLX313-firefly luciferase (pLX313-firefly luciferase was a gift from William Hahn and David Root Addgene plasmid # 118017; http://n2t.net/addgene:118017; RRID: Addgene_118017) plasmid using lentivirus production procedures described above. *In vitro* CAR T cytotoxicity assays were carried out as previously described^90^. Briefly, 5000 SCC8^luc^ or SCC42a^luc^ (antigen positive) cells were plated in 96-well plates in their respective culture mediums mixed 1:1 with ImmunoCult containing T cells at effector:target (E:T) ratios of 0.25:1, 0.5:1, 1:1, 2:1 and 4:1 in the presence of 0.75 µg/ml luciferin. Plates were read every 24 h and % specific lysis determined. *% specific lysis* = 100 * (*spontaneous lysis RLU – test RLU*)/(*spontaneous lysis RLU – kill control RLU*) where *spontaneous lysis* is defined as the no T cell control; *test* is defined as the well at the indicated E:T ratio and the *kill control* defined as the cells alone well treated with 2.5% NP-40 at assay endpoint.

#### Cytokine ELISA

500, 000 target cells and CAR T cells were co-cultured in 24-well plates at an E:T ratio of 1:1 for 72 h in triplicate. After 72 h, media and cells were collected and the cells pelleted at 300xg for 5 min. The supernatant was collected and assayed for cytokines by interpolation from a standard curve using the DuoSet human ELISA kit for IFN-γ (R&D Systems, DY285B) and human TNF-α (R&D Systems), according to manufacturer’s description.

### QUANTIFICATION AND STATISTICAL ANALYSIS

Where appropriate, the quantitative analyses (e.g. statistical tests used, exact value of n) can be found in the relevant methods sections and figure legends. Significance levels are stated in the main text and/or figures where they apply. Unless otherwise stated, data analyses and visualization were performed using R (v.4.1.2) with the following packages: *ggplot2* (v.3.5.1), *ggpubr* (v.0.6.0), *ggbeeswarm* (v.0.7.2), *ggrepel* (v.0.9.5), *ggsankey* (v.0.0.9999), *reshape2* (v.1.4.4), *ComplexHeatmap* (2.16.0), *ConsensusClusterPlus* (v.1.64.0), *plot3D* (v.1.4.1), *stringr* (v.1.5.1), *VennDiagram* (v.1.7.3) and *sqldf* (v.0.4.1.1).

## SUPPLEMENTARY FIGURE LEGENDS

**Figure S1.**
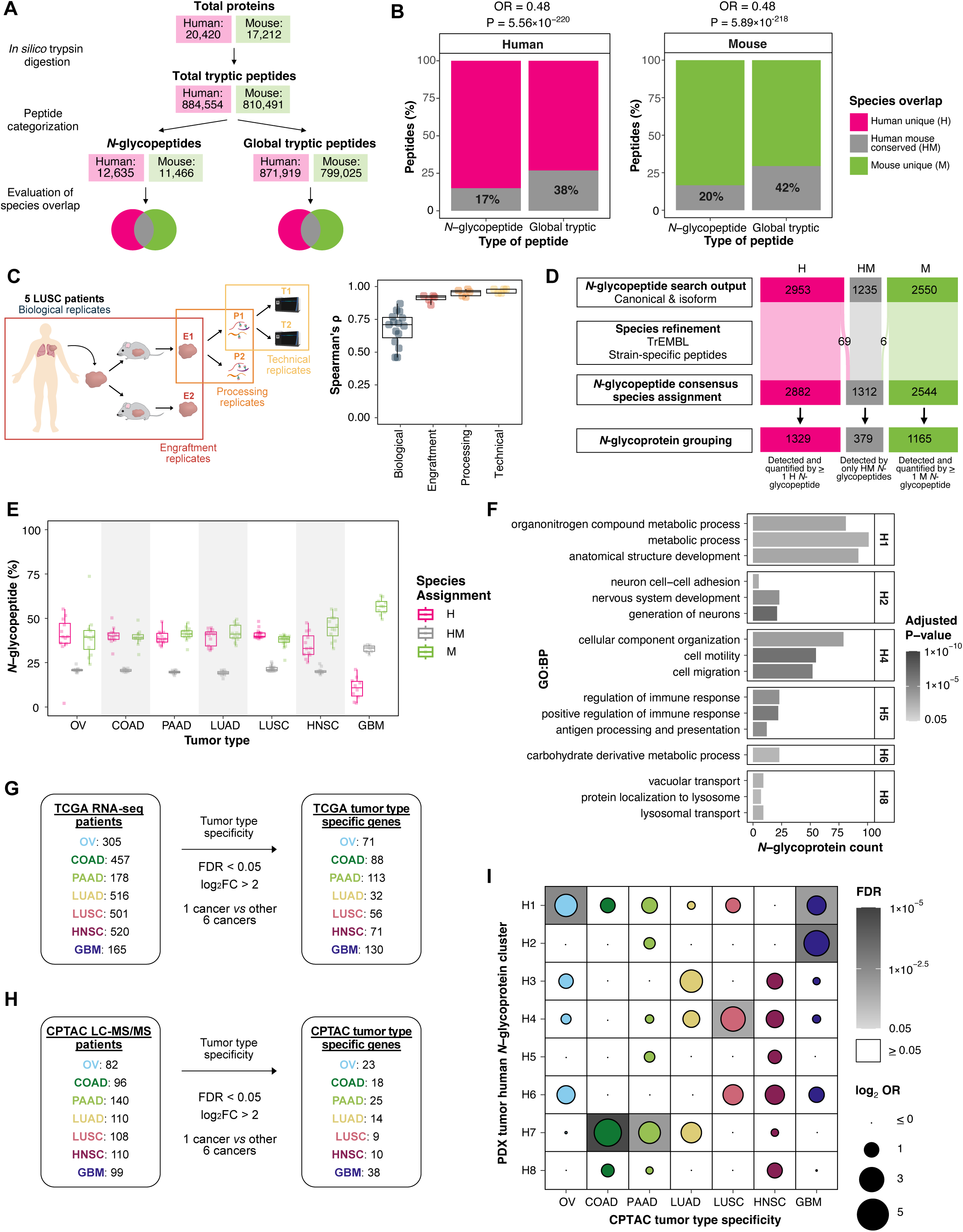
Pan-cancer PDX *N*-glycoproteome atlas of cancer-derived surface proteins, related to Figure 1. **(A)** Illustration of *in silico* human and mouse protein digest for species comparison. **(B)** Percentage of human-mouse conserved *N*-glycopeptides and global tryptic peptides in human (left) and mouse (right) proteomes. H = human unique, HM = human mouse conserved, M = mouse unique, OR= odds ratio. P-values calculated using Fisher’s exact test. **(C)** Schematic (left) and pairwise Spearman correlation analysis (right) of five PDX tumors for which engraftment, processing and technical replicates were analyzed. **(D)** Depiction of PDX *N*-glycopeptide species assignment and *N*-glycoprotein grouping. **(E)** Distribution of *N*-glycopeptides by species assignment per PDX tumor. **(F)** Select unique GO: Biological Processes enriched in human PDX *N*-glycoprotein clusters. No terms passed significance for H3 and H7 clusters. **(G & H)** Data analysis strategy to determine tumor type specific gene-products using **(G)** The Cancer Genome Atlas (TCGA) RNA-seq^17^ and **(H)** Clinical Proteomic Tumor Analysis Consortium (CPTAC) LC-MS/MS^18^ data. **(I)** Dot plot depicting over-representation of CPTAC tumor type specific proteins in respective PDX *N*-glycoprotein clusters. Dot size indicates the enrichment ORs, and background shading represents FDR.

**Figure S2:**
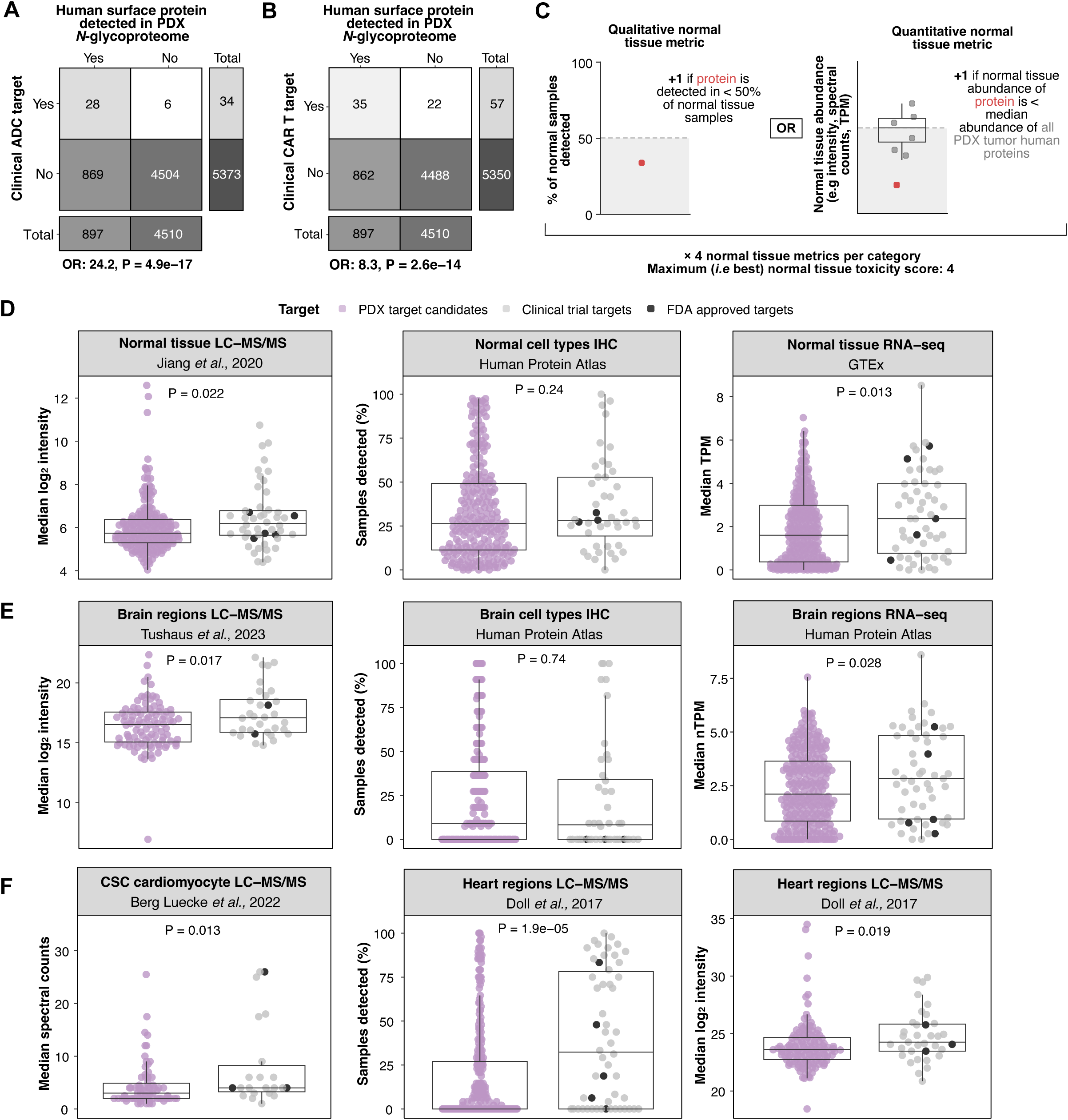
Prioritization and evaluation of cancer-enriched surface proteins with limited normal expression as therapeutic target candidates, related to Figure 2. **(A &B)** Contingency tables used to determine the likelihood of detecting a clinical **(A)** ADC and **(B)** CAR T target in the PDX human tumor *N*-glycoproteome against all other surface proteins in the human proteome. P-values were calculated using a Fisher’s exact test. **(C)** Normal tissue toxicity scoring system. **(D)** Distribution of full body normal tissue abundance or detection frequency of PDX target candidates (purple) *vs* solid tumor clinical immunotherapy targets (grey) in Jiang *et al.,*^20^ (left), Human Protein Atlas^5^ (middle) and GTEx RNA-seq data^19^ (right). **(E)** Distribution of normal brain tissue abundance or detection frequency of PDX target candidates (purple) *vs* solid tumor clinical immunotherapy targets (grey) in Tushaus *et al.,*^21^ (left), Human Protein Atlas brain IHC data^5^ (middle) and Human Protein Atlas brain RNA-seq data^22^ (right). **(F)** Distribution of normal heart tissue abundance or detection frequency of PDX target candidates (purple) *vs* solid tumor clinical immunotherapy targets (grey) in Berg Luecke *et al.,*^23^ (left) and Doll *et al.,*^24^ (middle & right). **(D-F)** P-values from unpaired Mann-Whitney U tests.

**Figure S3.**
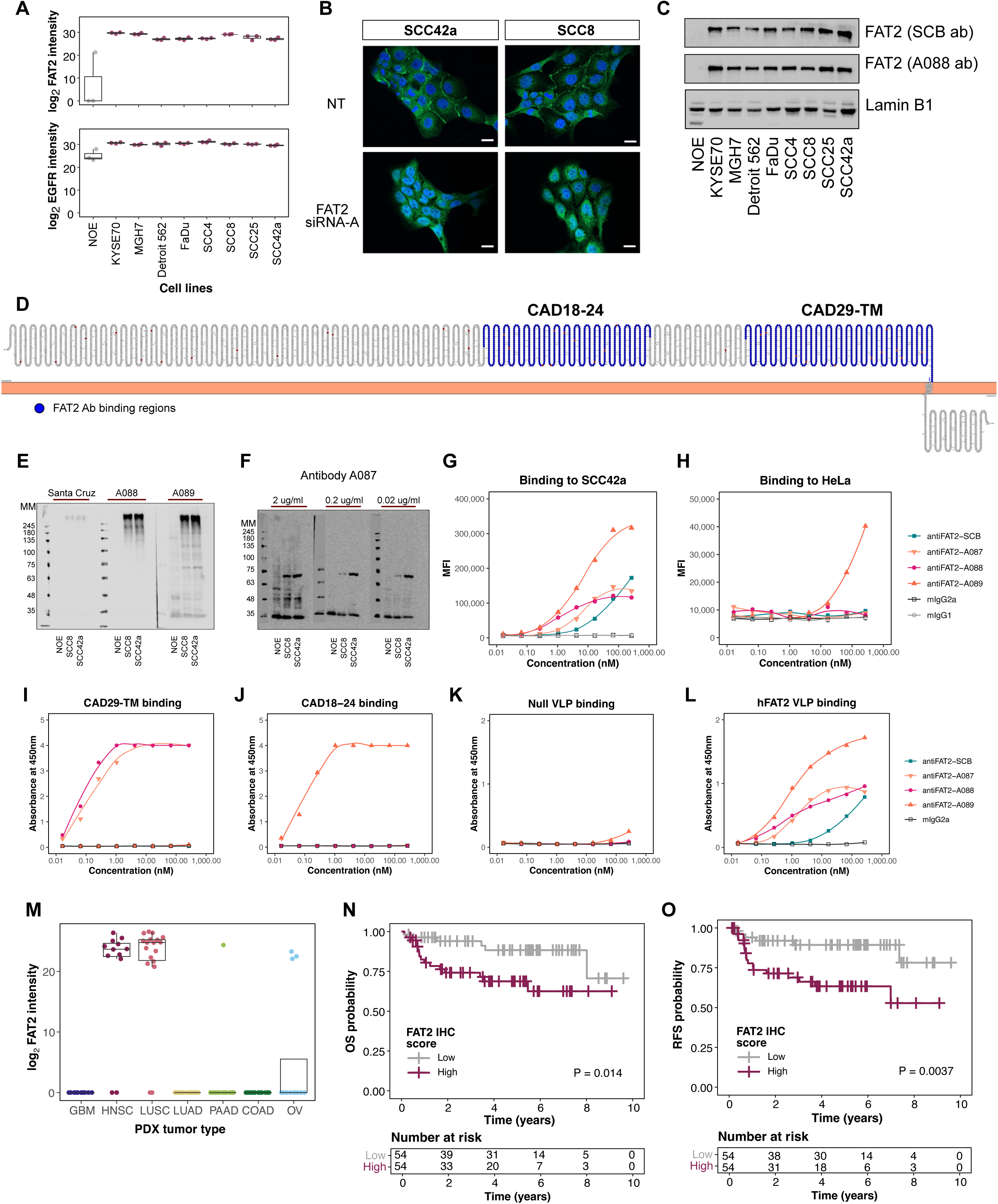
Identification and validation of FAT2 as a HNSC enriched surface protein, related to Figure 3. **(A)** Cell surface abundance of FAT2 (top) and EGFR (bottom) in NOE and SCC cell lines as determined by Cell Surface Capture (n=3). **(B)** Representative immunofluorescence microscopy images showing FAT2 expression on the surface of the HNSC cell lines SCC42a and SCC8. Top panel: Non-treated (NT) cells; bottom panel: siRNA knock-down of FAT2. Green: FAT2; Blue: DAPI; Scale bar: 20 µm. **(C)** Immunoblots performed with a commercial antibody (Santa Cruz Biotechnology [SCB] top panel) and a newly developed FAT2 antibody (A088) detect FAT2 expression in a panel of SCC cell lines. **(D)** Topology model of FAT2^91^. Indicated regions in blue (CAD18-24 and CAD29-TM) were used for immunization to generate novel FAT2 antibodies. **(E)** Immunoblot characterization of the newly developed anti-FAT2 clones A088 and A089 in the FAT2-negative cell line (NOE) and two FAT2-positive HNSC cell lines (SCC8, SCC42a). Results from commercial antibody (Santa Cruz) shown on the same gel. **(F)** Immunoblot characterization of the newly developed anti-FAT2 clone A087 in the FAT2-negative cell line (NOE) and two FAT2-positive HNSC cell lines (SCC8, SCC42a). **(G & H)** Binding assays of multiple anti-FAT2 candidate clones to compare the binding affinities to **(G)** SCC42a (FAT2-positive) and **(H)** HeLa cells (FAT2-negative). Two nonspecific isotype controls, mIgG2a, and mIgG1, were used. **(I-L)** Binding assays, performed with the ligands **(I)** CAD29-TM, **(J)** CAD18-24, **(K)** null virus-like particles (VLP), and **(L)** VLPs expressing full length FAT2, to test antibody specificities. **(M)** FAT2 abundance in our pan-cancer *N*-glycoproteomic PDX atlas. **(N & O)** HNSC patient **(N)** overall survival (OS) and **(O)** recurrence-free survival (RFS) probability based on median dichotomized FAT2 expression using the anti-FAT2 antibody A088. Statistical significance was calculated with a log-rank test.

**Figure S4.**
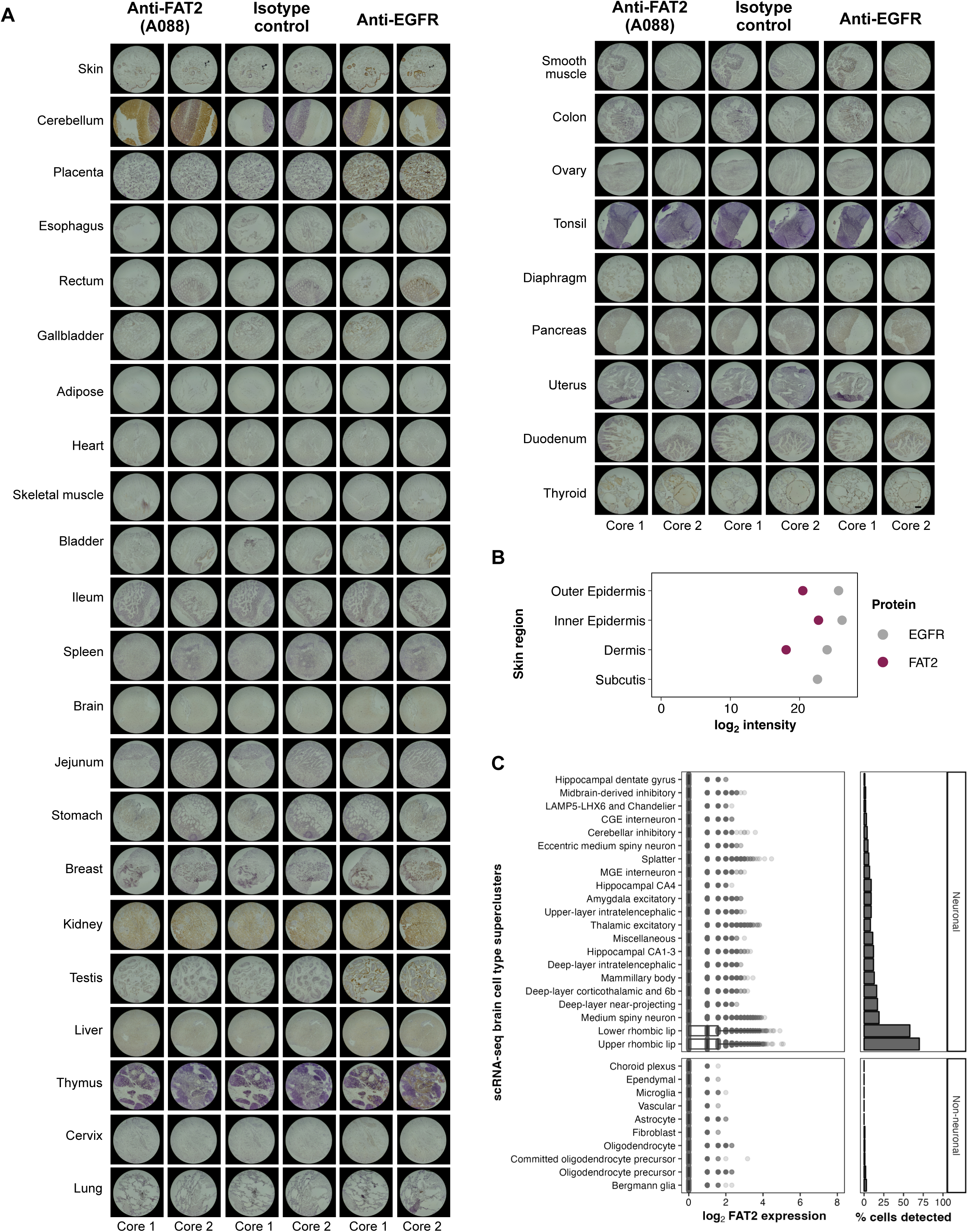
Validation of FAT2 as a surface protein with limited expression in normal tissue, related to Figure 3. **(A)** IHC staining of tissue cores available on a commercial normal tissue microarray (Novus Biologicals) using the FAT2 antibody A088. Isotype control (IgG2a) and epidermal growth factor receptor (EGFR) antibodies were used as negative and positive controls, respectively. Scale bar: 200 µm. **(B)** Proteomic abundance of FAT2 and EGFR in spatially resolved skin regions analyzed by Dyring-Andersen *et al.,*^34^. **(C)** *FAT2* expression (left) and frequency of detection (right) in normal brain cell type scRNA-seq clusters defined by Siletti *et al.,*^35^.

**Figure S5.**
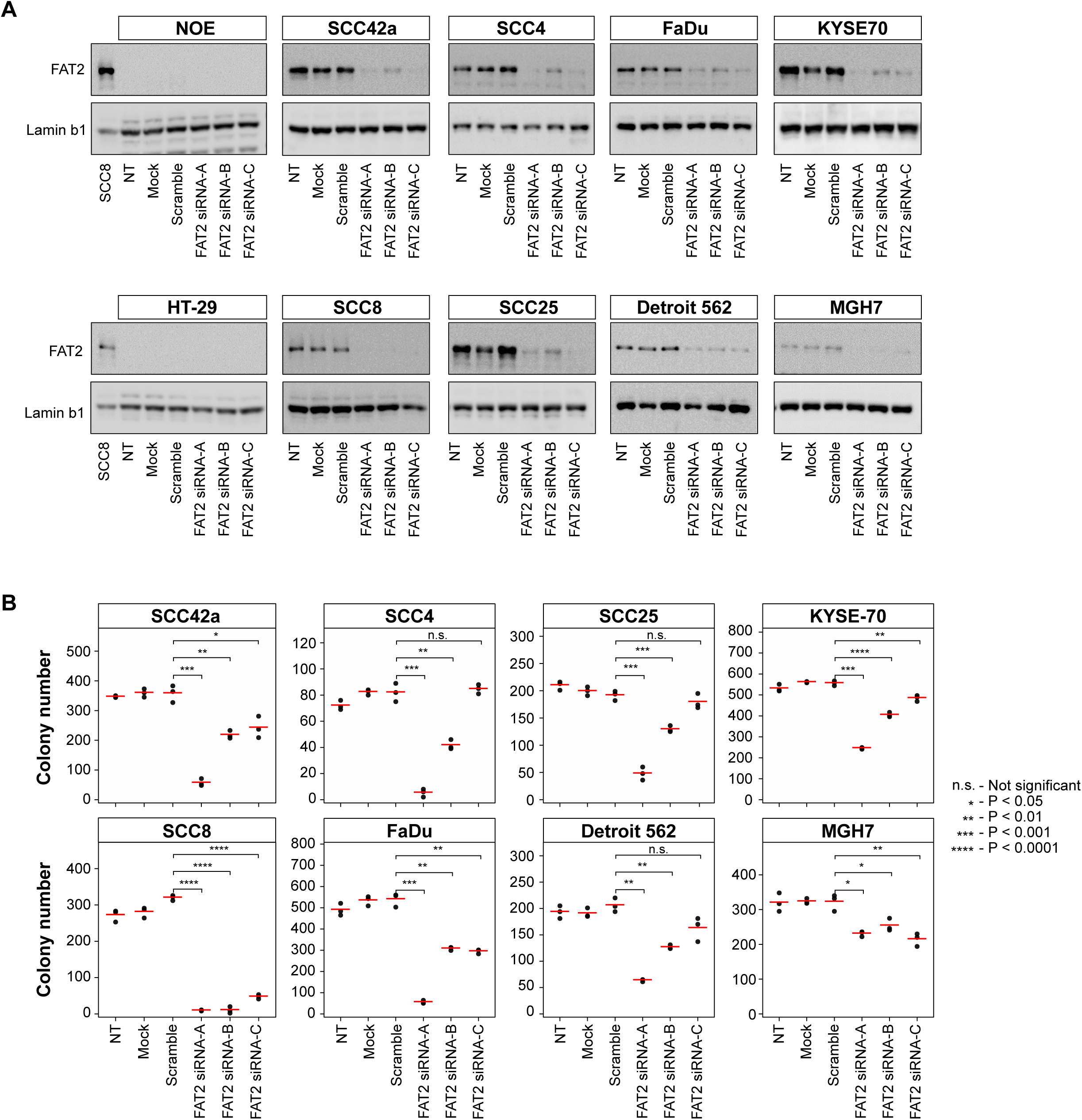
FAT2 is essential for HNSC growth and survival in vitro, related to Figure 4. **(A)** Immunoblot detection of FAT2 protein in two FAT2 negative cell lines (normal oral epithelial cells [NOE] and the colorectal adenocarcinoma cell line [HT-29]) and eight FAT2 positive SCC cell lines following siRNA-mediated knockdown. An SCC8 protein extract was used as a positive control in immunoblots using FAT2 negative cells. **(B)** Quantitation of colony numbers in SCC cell lines following siRNA-mediated knockdown (n = 3). P-values were determined with an unpaired Student’s t-test.

**Figure S6.**
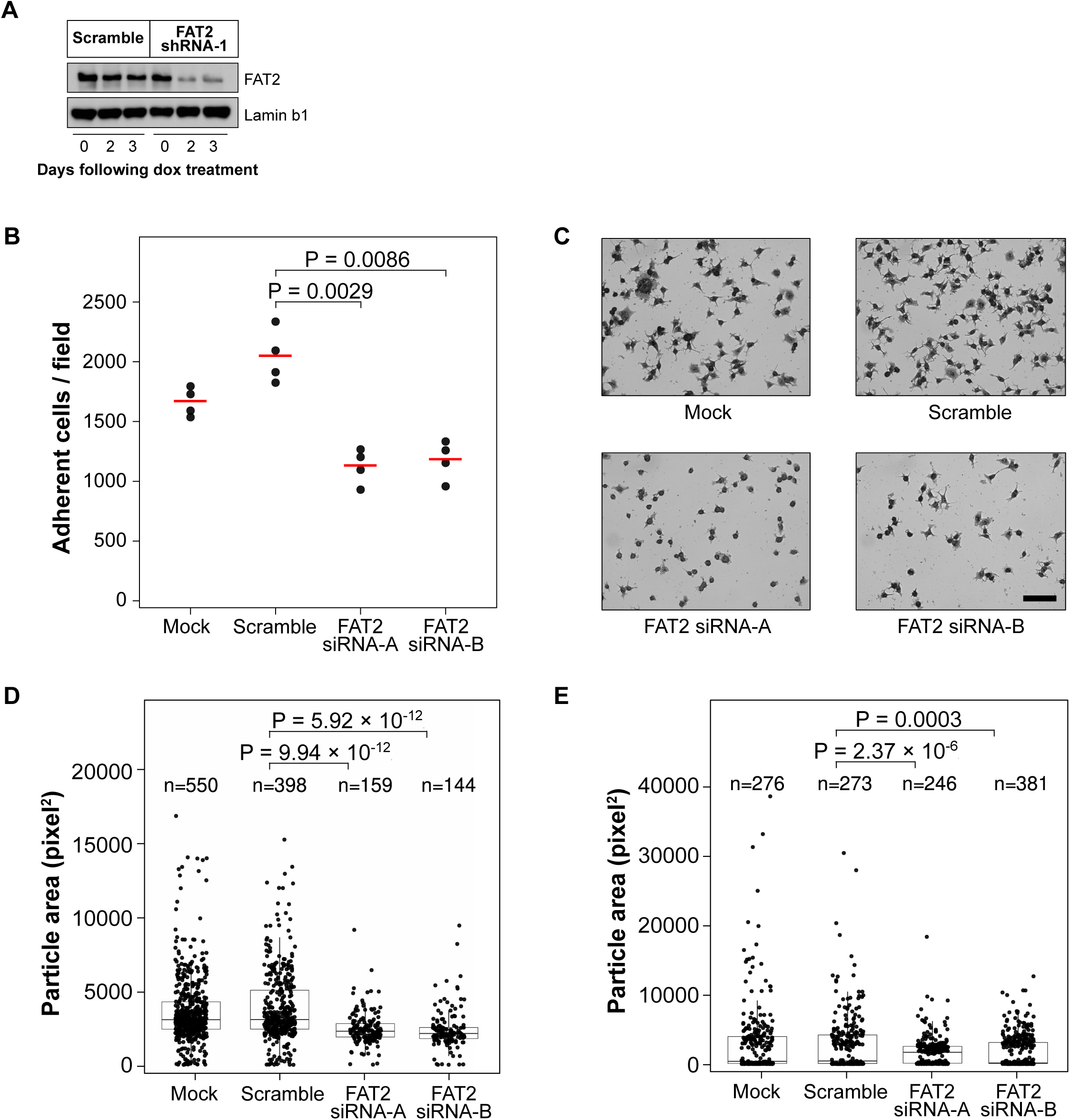
FAT2 mediated HNSC surface organization, adhesion and survival through integrin-PI3K signaling, related to Figure 6. **(A)** Immunoblot confirms shRNA-induced FAT2 depletion at timepoint used for Cell Surface Capture. **(B)** Quantitation of adherent cells to laminin matrix. Mean (red line) and individual data points (black) are visualized (n=4). P-values were calculated with an unpaired Student’s t-test. **(C)** Cell adhesion to laminin matrix of SCC42a cells following depletion of FAT2. Scale bar: 100 µm. **(D)** Cell size analysis of SCC42a cells following adhesion to collagen I, reported as particle area. Mean and quartile distribution shown. P-values were determined with an unpaired Student’s t-test. **(E)** Cell size analysis of SCC42a cells following adhesion to laminin matrix, reported as particle area. Mean and quartile distribution shown. P-values were determined with an unpaired Student’s t-test. All panels are representative of three independent experiments.

